# The immune response is a critical regulator of zebrafish retinal pigment epithelium regeneration

**DOI:** 10.1101/2020.08.14.250043

**Authors:** Lyndsay L. Leach, Nicholas J. Hanovice, Stephanie M. George, Ana E. Gabriel, Jeffrey M. Gross

**Affiliations:** Department of Ophthalmology, Louis J. Fox Center for Vision Restoration, University of Pittsburgh School of Medicine, Pittsburgh PA, 15213; Department of Biological Sciences, Kenneth P. Dietrich School of Arts and Sciences, University of Pittsburgh, Pittsburgh PA, 15260; Department of Developmental Biology, University of Pittsburgh School of Medicine, Pittsburgh PA, 15213

## Abstract

Loss of the retinal pigment epithelium (RPE) due to dysfunction or disease can lead to blindness in humans. Harnessing the intrinsic ability of the RPE to self-repair is an attractive therapeutic strategy; however, mammalian RPE is limited in its regenerative capacity. Zebrafish possess tremendous intrinsic regenerative potential in ocular tissues, including the RPE, but little is known about the mechanisms that drive RPE regeneration. Here, utilizing zebrafish, we identified elements of the immune response as critical mediators of intrinsic RPE regeneration. Macrophages/microglia are responsive to RPE damage and are required for the timely progression of the regenerative response and our data highlight that inflammation post-RPE injury is also critical for normal RPE regeneration. To our knowledge, these data are the first to identify the molecular and cellular underpinnings of RPE regeneration in any system and hold significant potential for translational approaches aimed towards promoting a pro-regenerative environment in mammalian RPE.

## INTRODUCTION

The retinal pigment epithelium (RPE) is a monolayer of polarized tissue that spans the entire posterior segment of the eye. Nestled between the light-sensing rod and cone photoreceptors of the neural retina and the nutrient-delivering blood vessels of the choroid vasculature, the RPE serves as an intermediary between these tissues and comprises the blood-retinal-barrier. As implicated by its location, the RPE performs myriad crucial functions to preserve the health and homeostasis of the surrounding retinal and vascular tissues; further, the RPE has a critical role in perpetuating the visual cycle and is integral to maintaining vision (1). Consequently, when the RPE becomes damaged or dysfunctional due to genetic mutation, injury, or disease, the functionality of the surrounding tissues is compromised, and vision is severely impaired. Primary insult to the RPE occurs in ocular degenerative diseases, including: Stargardt disease (STGD) (2–4); some forms of retinitis pigmentosa (RP) (5–7); and atrophic (or “dry”) age-related macular degeneration (AMD) (8, 9), which is the more common form of AMD and a leading cause of blindness worldwide (10). STGD and RP affect the very young (e.g. child to young adulthood onset (4, 7)), while AMD typically presents in individuals 50 years and older (10). There are currently no effective therapies for these RPE degenerative diseases, which presents an enormous healthcare burden, as 196 million people globally are projected to be living with AMD alone in 2020 (11). To compound the lack of treatment options, mammalian RPE and retinal tissues are limited in proliferative capacity, so tissue degeneration and consequent vision loss due to advanced disease in patients is irreversible. Gene therapy (12) and RPE cell-replacement therapeutics (13, 14) are currently in clinical trials, but an attractive alternate treatment option lies in harnessing the intrinsic regenerative capacity of the RPE.

Vertebrate retinal regeneration has been extensively studied in both amniotes (e.g. birds and mammals) and anamniotes (e.g. fish and frogs) (15–18); however, very little is known about the biology underlying vertebrate RPE regeneration. It is known that mammalian RPE can repair small lesions, but that larger-scale restoration is not possible or leads to over-proliferation and pathology (19). Some insight into the proliferative capacity of mammalian RPE has been gleaned from studies in mice (20, 21) and cultured human RPE cells (22), while studies in regeneration-capable non-mammalian systems have focused largely on RPE-to-retina transdifferentiation within the context of understanding retinal regeneration (23, 24). Thus, at present, the mechanisms driving intrinsic RPE regeneration remain elusive. Recently, our lab developed a zebrafish RPE ablation model and showed that the RPE has the intrinsic capacity to regenerate after widespread injury (25). This model uses a transgenic nitroreductase-mediated injury paradigm that enables tissue-specific ablation of the central two-thirds of the RPE. Extensive characterization showed: (i) sequential death of the RPE and photoreceptors, (ii) peripheral-to-central expansion of regenerating RPE, (iii) robust proliferation driving RPE regeneration, and (iv) recovery of visual function and RPE/photoreceptors post-injury (25).

Recent studies have converged on immune-related systems as required to resolve damage and stimulate regenerative responses in multiple model organisms and tissue contexts (e.g. (26–32)), including in the eye (e.g. (33–39)). Here, we utilized our zebrafish ablation model to identify the immune response as a critical mediator of RPE regeneration *in vivo*. Our data show that immune-related genes are upregulated in the RPE during early- and peak-stages of regeneration and that specific leukocytes respond to RPE ablation by infiltrating the injury site, undergoing changes in morphology, and expressing phagocytic and proliferative markers. Further, we show that RPE regeneration is impaired upon pharmacological dampening of inflammation and in an *irf8* mutant background, which is depleted of macrophages and lacks microglia (the macrophages of the central nervous system (CNS)) at larval stages (40). Collectively, the results of this study hold significant translational implications for mitigating RPE degenerative disease by revealing a role for the immune response in modulating the intrinsic ability of the RPE to regenerate.

## RESULTS

### Immune-related gene expression signatures are upregulated in RPE during regeneration

Our lab has shown that zebrafish have the capacity to regenerate RPE post-injury by utilizing a genetic ablation paradigm (25); however, the molecular components and signaling pathways involved in RPE regeneration remain unknown. This injury model utilizes a transgenic zebrafish line (*rpe65a*:nfsB-eGFP) wherein the RPE-specific (41) *rpe65a* enhancer drives expression of nitroreductase (nfsB) fused to eGFP for visualization and cell isolation. In the presence of nfsB, metronidazole (MTZ), a nontoxic prodrug, is converted into an apoptosis-inducing agent that results in targeted ablation of cells expressing the transgene (42). For all nfsB-MTZ ablation experiments, larvae were treated with 10mM MTZ for 24 hours, from 5-6 days post-fertilization (dpf; i.e. 0-1 days post-injury (dpi)), and subsequently allowed to recover. We utilized bulk RNA-sequencing (RNA-seq) as a global, unbiased approach to begin to identify RPE regenerative mechanisms. To isolate RPE, eyes were enucleated from unablated (MTZ-) and ablated (MTZ+) *rpe65a*:nfsB-eGFP larvae, dissociated into a single cell suspension, and eGFP^+^ RPE were isolated using fluorescence-activated cell sorting (FACS) for RNA-seq at three timepoints: 2dpi (7dpf age-matched controls); 4dpi (9dpf age-matched controls); and 7dpi (12dpf age-matched controls; Fig. 1A). These timepoints were chosen as prior characterization identified early- (2dpi), peak- (4dpi), and late-stages (7dpi) of RPE regeneration discernible by resolution of apoptosis, peak proliferation, and recovery of RPE marker expression, respectively (25). To determine genes and pathways upregulated during RPE regeneration, differential gene expression analyses were performed and enriched Reactome pathways were identified from filtered differentially expressed genes (DEGs; fold change ≥2, false discovery rate (FDR) ≤0.05). Innate/immune response- and complement-related gene sets were enriched at 4dpi (Fig. 1C); with cytokines, cytokine receptors, and matrix metalloproteases (MMPs) among the DEGs comprising these groups (Table S2). Similarly, at 2dpi, *mmp13b* and cytokine genes (e.g. *il11b, il34, cxcl8a, cxcl18b*) were upregulated when compared to 7dpf MTZ-controls; in fact, *il11b* was found to be the most highly upregulated gene at this early regenerative timepoint (Fig. 1D, Table S1). Importantly, RPE purification was confirmed using *rpe65a*, which was highly expressed in all eGFP^+^ cell populations, while leukocyte marker expression (e.g. *mpeg1.1* (43, 44), *lyz* (45), *mpx* (46)) was low or absent, indicating enrichment of RPE for RNA-seq, not leukocytes that have engulfed eGFP^+^ cell debris (Fig. S1). Leukocyte recruitment factors, *il34, cxcl8a*, and *cxcl18b*, were among the top-50 upregulated DEGs in RPE at both 2dpi and 4dpi (Tables S1 and S2). Interleukin-34 (IL-34) is a known ligand for the macrophage colony stimulating factor 1 receptor (CSF-1R; (47)) and an important signal for recruitment, survival, and differentiation of macrophages and microglia (48–50); while Cxcl8 (i.e. interleukin-8/IL-8) and Cxcl18b are potent neutrophil recruitment factors (51, 52). Differential expression of *il34, cxcl8a*, and *cxcl18b* was not significantly upregulated when comparing 7dpi and 12dpf datasets (Table S3), suggesting that leukocytes are no longer recruited at these late-stages post-RPE ablation. Lists of top downregulated DEGs can be found in Tables S4-S6. Consistent with previous literature highlighting a role for the immune response in multiple regenerative contexts, including the retina, these data indicate that immune-related genes, which include those encoding critical mediators of leukocyte recruitment and function, are also upregulated at early- and peak-stages during RPE regeneration.

**Figure 1.**
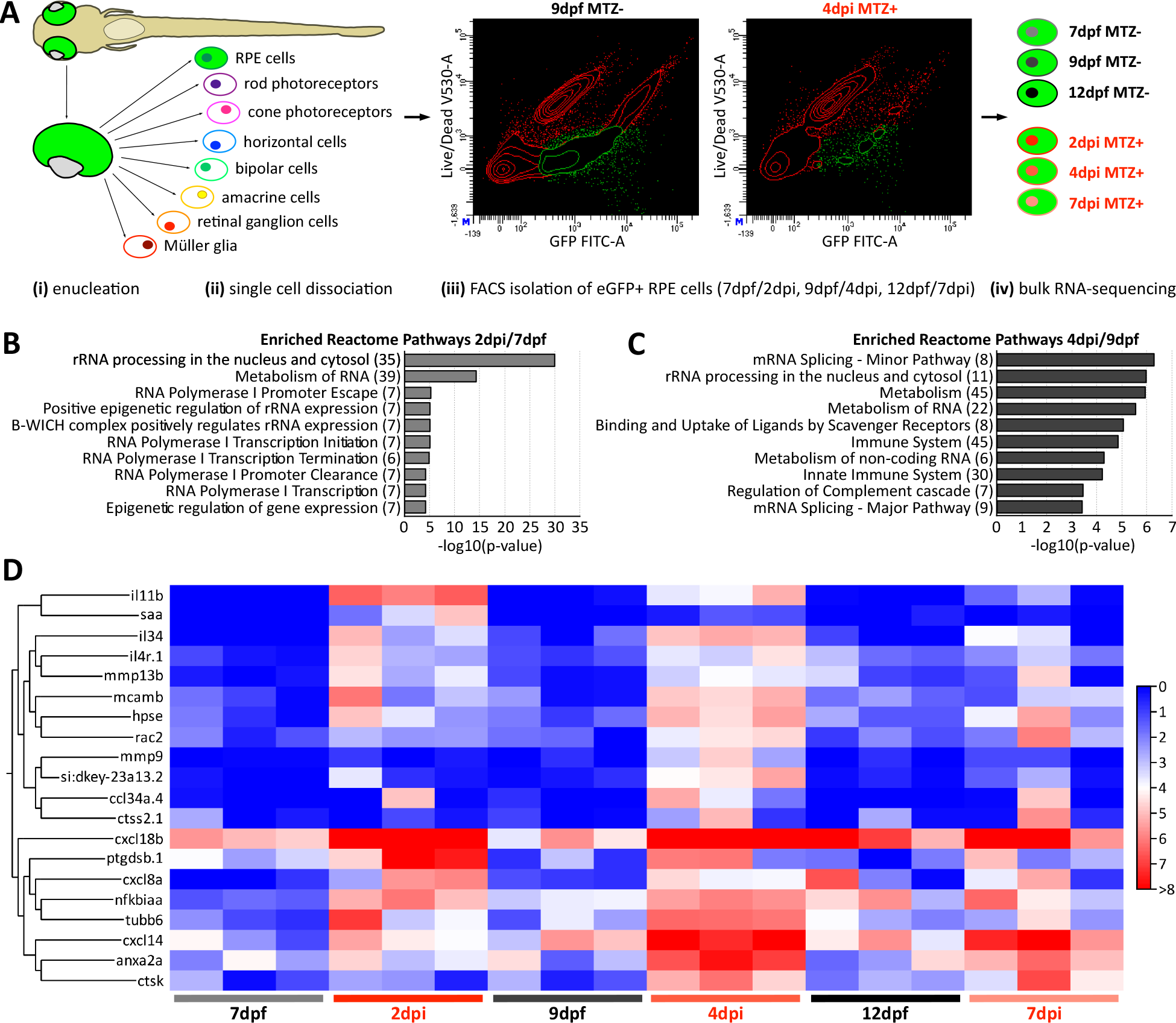
Enrichment of immune system genes during RPE regeneration. **(A)** Experimental workflow showing steps for tissue processing (*i,ii*) and isolation of eGFP^+^ RPE by FACS for bulk RNA-sequencing at different timepoints (*iii,iv*). Example 9dpf and 4dpi FACS plots show gate set for cell sorting (*iii; green*). **(B,C)** Enrichment analyses performed on groups of significant DEGs (fold change ≥2, FDR p-value ≤0.05) from 2dpi/7dpf **(B)** and 4dpi/9dpf **(C)** using the Reactome Pathway database. Data represent results from 332 genes from 3 experiments (2dpi/7dpf comparison) and 301 genes from 3 experiments (4dpi/9dpf comparison), and reveal enrichment of immune system, innate immune system, and complement-related gene sets at 4dpi. Numbers in parentheses indicate the number of significant DEGs enriched in each Reactome pathway. **(D)** Heatmap showing hierarchical clustering of immune-related genes in MTZ- and MTZ+ treatments across all timepoints. Genes were selected for representation based on presence in the top-50 upregulated gene sets from 2dpi/7dpf and 4dpi/9dpf DEG analyses (see Tables S1 and S2); this includes genes enriched in the Immune System and Innate Immune System Reactome Pathway groups **(C)**. Heatmap legend represents log_2_(TPM+1). Definitions as follows: DEGs, differentially expressed genes; dpf, days postfertilization; dpi, days post-injury; FACS, fluorescence-activated cell sorting; FDR, false discovery rate; MTZ, metronidazole; RPE, retinal pigment epithelium; TPM, transcripts per million.

### Macrophages/microglia respond post-RPE ablation at key regenerative timepoints

Next, we wanted to confirm specific leukocyte recruitment to the RPE injury site and understand the temporal dynamics of infiltration between the time spanning RPE-ablation and peak regeneration. Zebrafish do not develop adaptive immunity until at least 3 weeks post-fertilization (53), so only innate immune cells (neutrophils and macrophages) were examined for these experiments. Neutrophil infiltration was assessed in unablated (MTZ-) and ablated (MTZ+) larvae using the transgenic reporter line, *lyz*:TagRFP (45). In the majority of unablated larvae, there were no *lyz*:TagRFP^+^ cells observed in the RPE in whole mount eyes from 6-9dpf (Fig. 2A-D). In ablated larvae, few *lyz*:TagRFP^+^ cells were apparent in the RPE from 1-4dpi (Fig. 2E-H; *white arrowheads*). Quantification of the number of cells showed no significant difference in infiltration at 1dpi, 3dpi, or 4dpi, but a significant increase in the number of *lyz*:TagRFP^+^ cells at 2dpi when compared to 7dpf controls (Fig. 2Q; p=0.0038). However, while the results at 2dpi were significant, six neutrophils was the maximum number observed in the RPE of any larva, and this was only in one larva from this dataset (Fig. 2F,Q).

**Figure 2.**
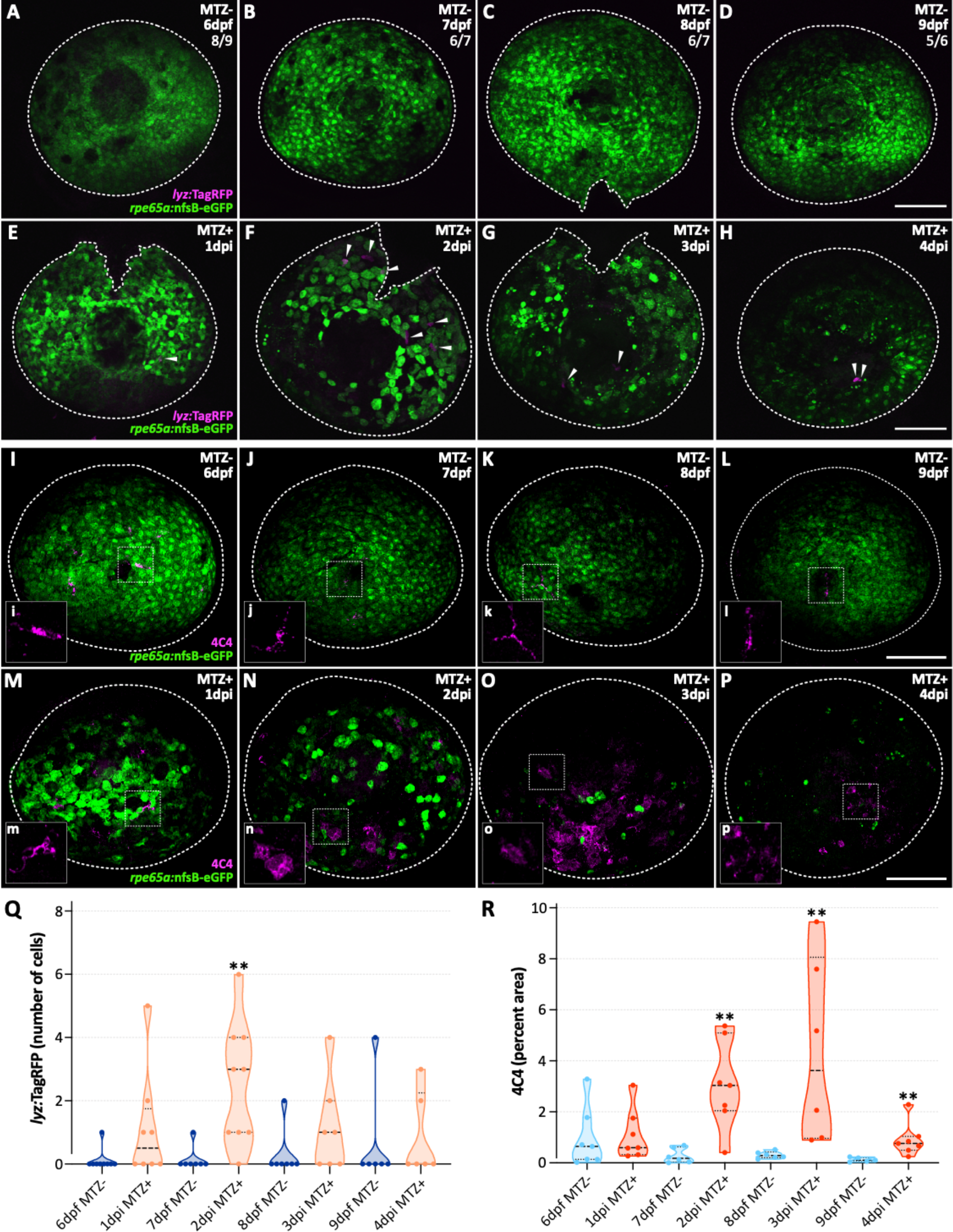
Leukocyte infiltration into the RPE injury site during regeneration. Fluorescent confocal micrographs of MTZ-unablated **(A-D)** and MTZ+ ablated **(E-H)** Tg(*lyz*:TagRFP; *rpe65a*:nfsB-eGFP) whole mount eyes from 6dpf/1dpi to 9dpf/4dpi. Images show only a few *lyz*:TagRFP^+^ neutrophils infiltrate the RPE post-ablation. **(A-D)** Ratios at top right indicate number of larval eyes lacking *lyz*:TagRFP^+^ neutrophils over total number of eyes for that timepoint (showing representative images). **(E-F)** *White arrowheads* point to *lyz*:TagRFP^+^ neutrophils. **(I-P)** Fluorescent confocal micrographs of MTZ-unablated **(I-L)** and MTZ+ ablated **(M-P)** Tg(*rpe65a*:nfsB-eGFP) whole mount eyes from 6dpf/1dpi to 9dpf/4dpi labeled with 4C4 antibody to mark MΦs/μglia. **(i-p)** Insets show digital zooms of single cells or clusters of cells to highlight 4C4^+^ cell morphologies. Images show resident 4C4^+^ microglia with ramified morphology in unablated larvae and an influx of 4C4^+^ MΦs/μglia post-RPE ablation; infiltrating cells appear in clusters with a more rounded morphology. **(A-P)** *White dashed lines* designate edge of whole mount larval eye. *Magenta* labels either endogenous *lyz*:TagRFP or 4C4 and *green* labels endogenous *rpe65a*:nfsB-eGFP. Scale bars represent 100μm. **(Q)** Violin plots showing quantification of the number of *lyz*:TagRFP^+^ neutrophils infiltrating during RPE regeneration. Data show a significant increase in number of neutrophils at 2dpi (MTZ+) when compared to 7dpf (MTZ-) controls (p=0.0038). A maximum of 6 infiltrating cells were present at the 2dpi (MTZ+) timepoint (datapoint shown in **F**). **(R)** Violin plots showing quantification of the percent area occupied by 4C4^+^ staining during RPE regeneration. Data show significant increases in MTZ+ ablated larvae at 2dpi (p=0.0047), 3dpi (p=0.0022), and 4dpi (p=0.0025) when compared to age-matched MTZ-unablated controls. **(Q,R)** *Dashed black lines* represent the median and *dotted black lines* represent quartiles. Statistics (number of experiments, biological replicates (n), statistical test, and p-values) can be found in Table 2. Dorsal is up; definitions as follows: **, p-value ≤0.01; dpf, days postfertilization; dpi, days post-injury; MΦs/μglia, macrophages/microglia; MTZ, metronidazole; RPE, retinal pigment epithelium.

To assess the dynamics of macrophage infiltration, both a 4C4 antibody (Fig. 2I-P,R; (54)) and the transgenic reporter line, *mpeg1*:mCherry (Fig. S2; (44)), were used for visualization. Notably, neither method distinguishes between monocyte-derived macrophages recruited post-injury and tissue resident macrophage (or microglia) populations, so cells from ablation studies will be referred to collectively as macrophages/microglia (MΦs/μglia) hereafter. Resident 4C4^+^ and mCherry^+^ MΦs/μglia were present in the RPE of unablated larvae in whole mount (Fig. 2I-L) and sectioned tissue (Fig. S2A,B,E,F,I,J,M,N; *white arrowheads*), respectively. From 2-4dpi, ablated larvae showed a visible increase in 4C4 (Fig. 2N-P) and mCherry signal (Fig. S2G,H,K,L,O,P; *white arrowheads and dotted brackets*). Due to the morphology and apparent clustering of both 4C4^+^ and mCherry^+^ cells, percent area quantification was performed instead of cell counts as individual cells were difficult to distinguish (e.g. Fig. 2O). Quantification of the percent area of 4C4 signal revealed a significant increase from 2-4dpi, with peak infiltration occurring at 3dpi (Fig. 2O,R). Similarly, percent area quantification of the mCherry signal showed a significant increase from 2-4dpi and a peak at 3dpi (Fig. S2K,L,Q). Importantly, MTZ treatment alone did not mobilize MΦs/μglia to the RPE as influx of mCherry^+^ cells was not observed at 4dpi in larvae lacking the *rpe65a*:nfsB-eGFP transgene (Fig. S2R).

In addition to an increased presence of MΦs/μglia post-RPE ablation, we also noted phenotypic differences in 4C4^+^ cells between unablated and ablated larvae (Fig. 2I-P; *insets*). Macrophages and microglia are highly dynamic cells that undergo a range of morphological changes both at rest and in response to injury and disease; this process is complex and controversial, but an overly simplified summary (from studies in the brain) is that ramified cells retract cellular extensions and become rounded (or “amoeboid”) after injury, which may signify phagocytic function (55, 56). Unablated larvae age 6-9dpf showed 4C4^+^ cells with extended branches and what appeared to be ramified morphology (Fig. 2i-l; *insets*), while ablated larvae showed more amoeboid 4C4^+^ cells, which was most obvious at the 2-4dpi timepoints (Fig. 2m-p; *insets*). These changes led us to look more closely at cell morphology, in particular, the rounding up of cells, from 2-4dpi, the times when MΦs/μglia infiltrate the RPE. To assess MΦ/μglia morphology *in vivo* and in three-dimensional space (3D), we utilized light-sheet microscopy and the transgenic reporter line, *mpeg1.1*:Dendra2 (57). Dendra2 is a photo-convertible protein that undergoes an irreversible switch from green to red fluorescence when exposed to violet or ultraviolet light (58); here, conversions were performed immediately prior to imaging for all larvae. We utilized Imaris to 3D-render, isosurface, and quantify the sphericity (59) of anterior *mpeg1.1*:Dendra2 (red)^+^ cells in ablated and unablated larvae from 2-4dpi (Fig. 3, Fig. S3). Sphericity measurements revealed that at 3dpi, but not 2dpi and 4dpi (Fig. S3C,F; p=0.8939 and p=0.2427, respectively), ablated larvae had *mpeg1.1*:Dendra2 (red)^+^ cells that were significantly more spherical, when compared to 8dpf unablated siblings (Fig. 3C; p=0.0298). These data suggest that MΦ/μglia may be phagocytic and responding to RPE injury during the time of peak infiltration.

**Figure 3.**
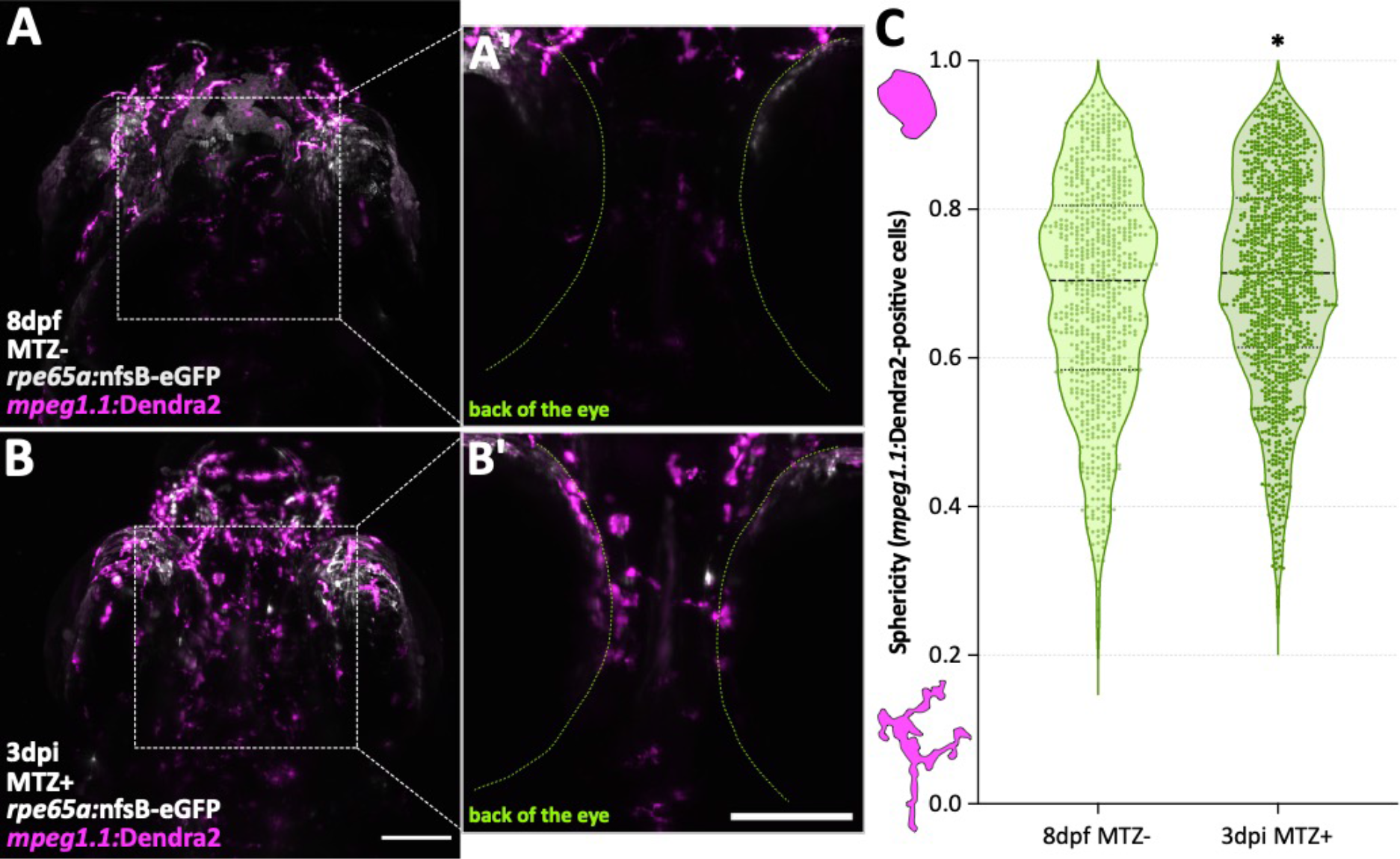
Anterior macrophages/microglia show increased sphericity at 3 days post-RPE ablation. *In vivo* fluorescent light-sheet micrographs from 8dpf MTZ-unablated **(A,A’)** and 3dpi MTZ+ ablated **(B,B’)** Tg(*mpeg1.1*:Dendra2; *rpe65a*:nfsB-eGFP) whole larvae (representative ~300μm z-stacks). **(A’,B’)** Digital zooms of cropped 100μm z-stacks (z-step 100-200) showing *mpeg1.1*:Dendra2 (red)^+^ cell localization in and adjacent to the back of the eye (*green dashed line*) in MTZ-unablated **(A’)** and MTZ+ ablated siblings **(B’)**. **(A-B’)** *White* labels endogenous *rpe65a*:nfsB-eGFP and *magenta* labels endogenous photoconverted *mpeg1.1*:Dendra2. Scale bars represent 100μm. **(C)** Violin plots showing quantification of the sphericity of *mpeg1.1*:Dendra2 (red)^+^ cells in the anterior portion of zebrafish larvae. Cells from 3dpi MTZ+ ablated larvae are significantly more spherical (rounded) than cells from MTZ-controls (p=0.0298). Statistics (number of experiments, biological replicates (n), statistical test, and p-values) can be found in Table 2. Data represent n=753 cells from 3 larvae (MTZ-) and n=1165 cells from 3 larvae (MTZ+). *Dashed black lines* represent the median and *dotted black lines* represent quartiles. Anterior is up; definitions as follows: *, p-value ≤0.05; dpf, days post-fertilization; dpi, days post-injury; MTZ, metronidazole; RPE, retinal pigment epithelium.

To directly assess MΦ/μglia gene expression during RPE regeneration, we utilized *mpeg1*:mCherry zebrafish (44) and performed bulk RNA-seq analyses on FACS-isolated mCherry-expressing MΦs/μglia at 2 and 4 days post-ablation using a similar workflow as for RPE RNA-seq (Fig. S4A-D). The same early- (2dpi) and peak- (4dpi) regeneration stages were selected to align with the RPE RNA-seq timepoints; however, we chose to forego analysis at 7dpi as there were few DEGs in RPE at this timepoint (Tables S3 and S6) and MΦ/μglia infiltration appeared to wane after 3dpi (Fig. 2R, Fig. S2Q). Pathway analysis of significant upregulated DEGs (fold change ≥2, FDR ≤0.05) showed that cell cycle- and mitosis-related Reactome gene sets were among the top-20 highly-enriched pathways in the MΦs/μglia of ablated larvae at 2dpi and 4dpi when compared to age-matched sibling controls (Fig. 4A,B). Full lists of enriched Reactome pathways at each timepoint can be found in Tables S9 and S10. Genes within the top-enriched Reactome Cell Cycle, Mitotic pathway were also found to be among the top-50 most significantly upregulated DEGs at both timepoints, and these included *cdk1*, cyclins, cell division cycle (cdc), and cytokinesis genes, among others (Fig. 4C, Tables S7 and S8). As with the RPE RNA-seq dataset, purification of MΦs/μglia was confirmed using *mpeg1.1*, which was highly expressed in all mCherry^+^ cell populations, while neutrophil (e.g. *lyz* and *mpx*) and RPE (*rpe65a*) marker expression was low or absent in all but one 7dpf replicate (Fig. S4E). It is known that macrophage populations proliferate in response to injury, inflammatory, and other disease stimuli (60) and that IL-34 is a pro-proliferative factor for these cells *in vivo* and *in vitro* (50). We have shown that isolated RPE express *il34* at 2dpi and 4dpi (Fig. 1D, Tables S1 and S2), suggesting that MΦs/μglia may be proliferating in response to RPE-derived cytokine stimulus. Furthermore, we detected significant upregulation of annexin A1, *anxa1a*, expression at 2dpi (Table S7). AnxA1 has been associated with macrophage phagocytosis (61, 62) and expression at this timepoint supports the observed morphology changes in 4C4^+^ cells at 2dpi (Fig. 2N) and indicates that infiltrating cells may be rounding up to clear debris.

**Figure 4.**
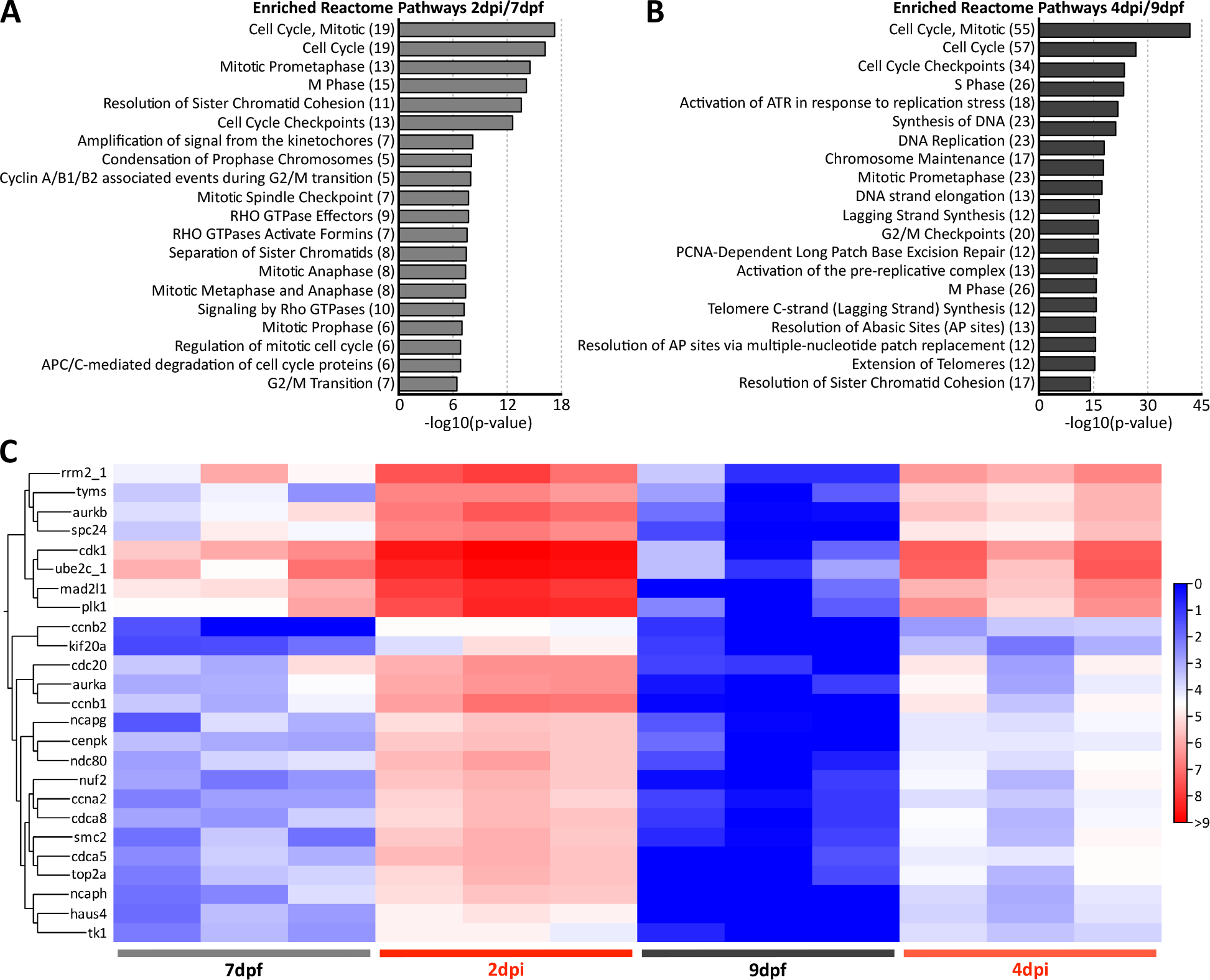
Cell cycle genes are upregulated in macrophages/microglia during RPE regeneration. **(A,B)** Top-20 Reactome pathways enriched from groups of significant DEGs (fold change ≥2, FDR p-value ≤0.05) at 2dpi/7dpf (**A**; 67 genes from 3 experiments) and 4dpi/9dpf (**B**; 208 genes from 3 experiments) in FACS-isolated mCherry^+^ MΦs/μglia. Data reveal enrichment of cell cycle- and mitosis-related gene sets at 2dpi and 4dpi. Numbers in parentheses indicate the number of significant DEGs enriched in each Reactome pathway. **(C)** Heatmap showing hierarchical clustering of cell cycle- and mitosis-related genes in MTZ- and MTZ+ treatments across both timepoints. Genes were selected for representation based on presence in the top-50 upregulated gene sets from 2dpi/7dpf and 4dpi/9dpf DEG analyses (Tables S7 and S8); this includes genes enriched in the Cell Cycle, Mitotic Reactome pathway group **(C)**. Heatmap legend represents log_2_(TPM+1). Definitions as follows: DEGs, differentially expressed genes; dpf, days post-fertilization; dpi, days post-injury; FACS, fluorescence-activated cell sorting; FDR, false discovery rate; MΦs/μglia, macrophages/microglia; MTZ, metronidazole; RPE, retinal pigment epithelium; TPM, transcripts per million.

Collectively, these data show that MΦs/μglia are the responsive leukocyte after RPE injury and during RPE regeneration. MΦ/μglia infiltration peaked at 3dpi and, at this timepoint, cells may be responding to RPE injury by taking on a more amoeboid morphology. RNA-seq revealed that MΦs/μglia are proliferating in ablated larvae at 2dpi and 4dpi, which is when *il34* (a pro-proliferative signal) is significantly upregulated in regenerating RPE. MΦ/μglia cells play dual roles in both promoting and resolving inflammation and have the ability to switch from pro- to anti-inflammatory phenotypes to restore damaged tissue (63); thus, these findings prompted us to more closely investigate both inflammation, and MΦs/μglia specifically, as critical mediators of RPE regeneration.

### Pharmacological inhibition of inflammation impairs zebrafish RPE regeneration

First, we wanted to determine if broadly dampening inflammation affected RPE regeneration in larval zebrafish. Larvae were exposed to 50μM dexamethasone, a synthetic glucocorticoid (GC) and potent anti-inflammatory agent, or 0.05% dimethyl sulfoxide (DMSO; vehicle control) for 24-hours prior to RPE ablation, for the 24-hour duration of ablation with MTZ, and for the entirety of the regeneration period, which was taken out to 4dpi (Fig. 5A). Efficacy of systemic dexamethasone treatment was confirmed by quantifying expression of the *pregnane X receptor* (*pxr*), a known transcriptional target of dexamethasone (64–66), which was found to be upregulated after a 24-hour dexamethasone exposure (Fig. 5B). Previously, proliferation and pigment recovery were quantified as metrics of the extent of RPE regeneration post-ablation (25). Other studies in the retina have used a similar approach, assaying proliferation and cell recovery as a readout for repair after treatment with dexamethasone (35, 67). As proliferation within the regenerating RPE peaks between 3-4dpi and visible pigment is largely recovered by 4dpi (25), we used the same approach here. Proliferation was assayed by treating larvae with 10mM BrdU for 24 hours, from 3-4dpi (Fig. 5A). Incorporation results showed significantly fewer RPE-localized BrdU^+^ cells in dexamethasone-treated 4dpi larvae when compared to 4dpi DMSO-treated siblings (Fig. 5C,E; *white arrowheads*; Fig. 5G; p=0.0002). Dexamethasone treatment showed no effect on cell proliferation in 9dpf unablated siblings (Fig. S5B-D; p=0.7579). Pigment recovery, which was quantified based on central expansion of continuous pigmented tissue (Fig. S5A), also lagged in dexamethasone-treated 4dpi ablated larvae (Fig. 5D,F; *magenta arrowheads*) and was significantly decreased compared to DMSO-treated controls (Fig. 5H; p=0.0262). To determine if dexamethasone impaired MΦ/μglia recruitment, larvae were aged to 8dpf/3dpi (peak MΦ/μglia infiltration) and stained with 4C4 antibody. As anticipated, there were few 4C4^+^ cells in 8dpf unablated controls (Fig. S6A,B) and abundant 4C4^+^ signal in 3dpi ablated DMSO controls (Fig. S6C). Interestingly, 4C4 also labeled dexamethasone-treated 3dpi ablated larvae (Fig. S6D) and there was no significant difference between these larvae and DMSO-treated controls (Fig. S6E; p=0.1143). Together, these data indicate that inflammation is necessary for RPE regeneration, but dexamethasone-mediated impairment of inflammation does not impact recruitment of leukocytes post-ablation.

**Figure 5.**
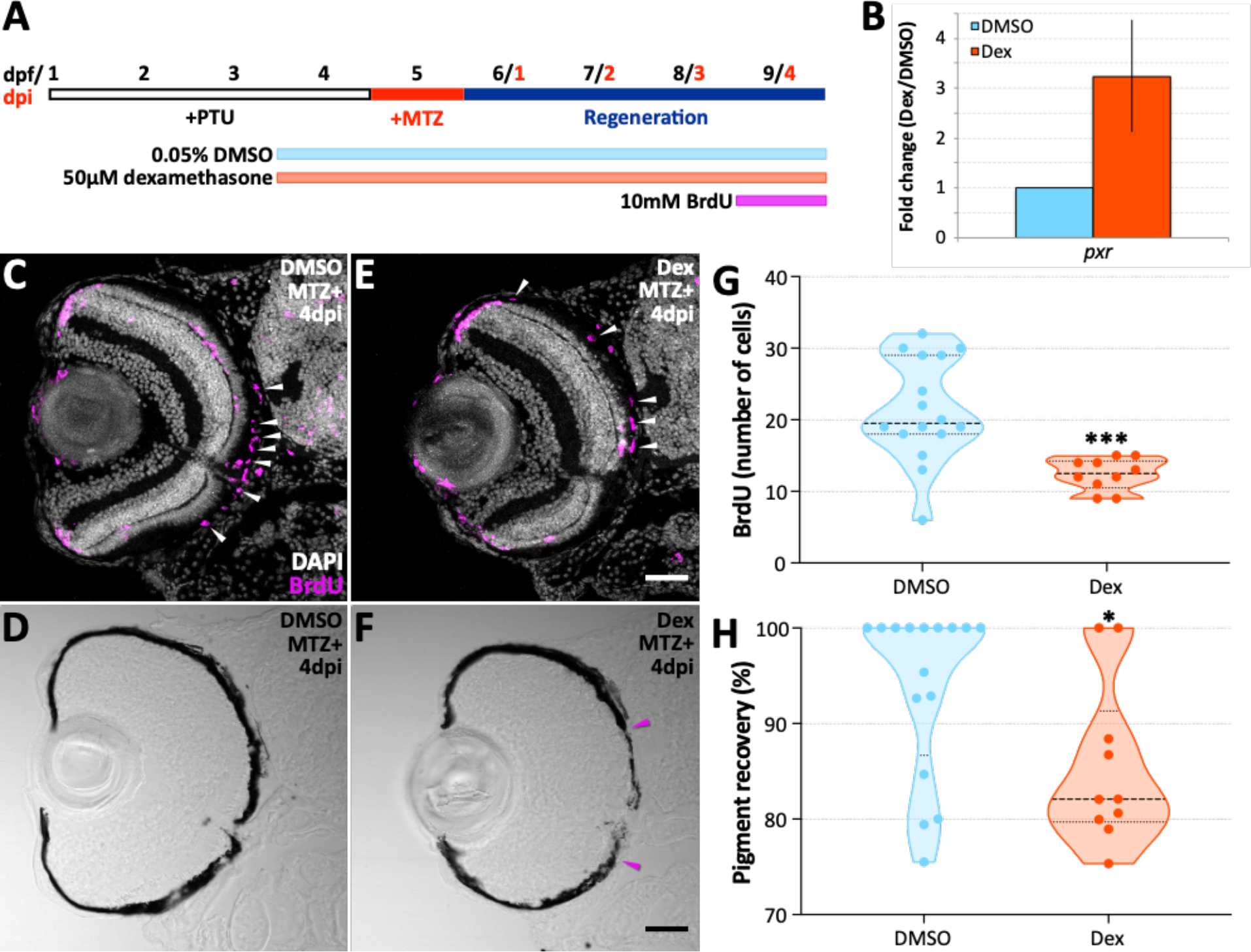
Suppression of inflammation with dexamethasone impairs RPE regeneration. **(A)** Schematic depicting the timeline of experimental, control, and BrdU labeling treatments out to 9dpf/4dpi. **(B)** Bar graph showing fold change in *pxr* gene expression by qRT-PCR analyses from larvae treated with dexamethasone (n=74 larvae, 3 separate experiments) or DMSO (n=71 larvae, 3 separate experiments) for 24 hours (4-5dpf). Error bars represent 95% confidence interval; *βactin* was used as a reference gene. **(C-F)** Fluorescent and brightfield confocal micrographs of transverse sections from 4dpi MTZ+ ablated DMSO- **(C,D)** and dexamethasone-treated **(E,F)** Tg(*rpe65a*:nfsB-eGFP) larval eyes. Images show central localization of BrdU-labeling and continuous pigmentation in DMSO-treated larvae, whereas dexamethasone-treated larvae show fewer BrdU^+^ cells and a gap in centralized pigmentation. **(C,E)** *White arrowheads* highlight BrdU-labeled cells (*magenta*) in the RPE layer. *White* (DAPI) labels nuclei. **(F)** *Magenta arrowheads* designate the edges of a regenerating RPE monolayer in dexamethasone-treated larvae. Scale bars represent 40μm. **(G,H)** Violin plots showing a significant decrease in the number of BrdU^+^ cells **(G)** and the percent recovery of a pigmented monolayer **(H)** in dexamethasone-treated larvae when compared to DMSO controls (p=0.0002 and p=0.0262, respectively). *Dashed black lines* represent the median and *dotted black lines* represent quartiles. Statistics (number of experiments, biological replicates (n), statistical test, and p-values) can be found in Table 2. Dorsal is up; definitions as follows: *, p-value ≤0.05; ***, p-value ≤0.001; BrdU, bromodeoxyuridine; dex, dexamethasone; dpf, days post-fertilization; dpi, days post-injury; DMSO, dimethyl sulfoxide; MTZ, metronidazole; PTU, n-phenylthiourea; *pxr*, pregnane X receptor; RPE, retinal pigment epithelium.

### Macrophages/microglia are required for zebrafish RPE regeneration

Next, we wanted to determine if MΦ/μglia cells, specifically, are required for RPE regeneration by quantifying proliferation and RPE recovery post-ablation in an *interferon regulatory factor 8* (*irf8*) mutant zebrafish line (40). Irf8 is an important regulator of monocyte/macrophage (68) and microglia (69) lineages. *irf8* mutant zebrafish lack macrophages during the early stages of larval development, which begin to recover by 7dpf, although these cells remain immature (40). Additionally, *irf8* mutants are completely devoid of microglia to 31dpf (40). To assess proliferation post-RPE ablation, *irf8* wild-type and mutant larvae were incubated in 10mM BrdU from 3-4dpi and fixed immediately thereafter for analyses. At 4dpi, *irf8* mutants unexpectedly showed a significant increase in numbers of BrdU^+^ cells in the RPE when compared to wild-type siblings (Fig. 6A-C; p=0.0111). There was no significant difference in BrdU incorporation between age-matched unablated (MTZ-) *irf8* wild-type and mutant controls (Fig. 6C; p=0.1054), indicating retention of proliferating cells was a result of MTZ-dependent RPE ablation and not the *irf8* mutation itself. While increased proliferation was unanticipated based on our dexamethasone treatment results (Fig. 5G), we also noted substantial accumulation of pyknotic nuclei between the photoreceptor layer and the RPE only in ablated *irf8* mutant larvae (Fig. 6B,B’; *white dotted lines*). A similar phenotype has been noted in *irf8* mutants post-spinal cord injury (27).

**Figure 6.**
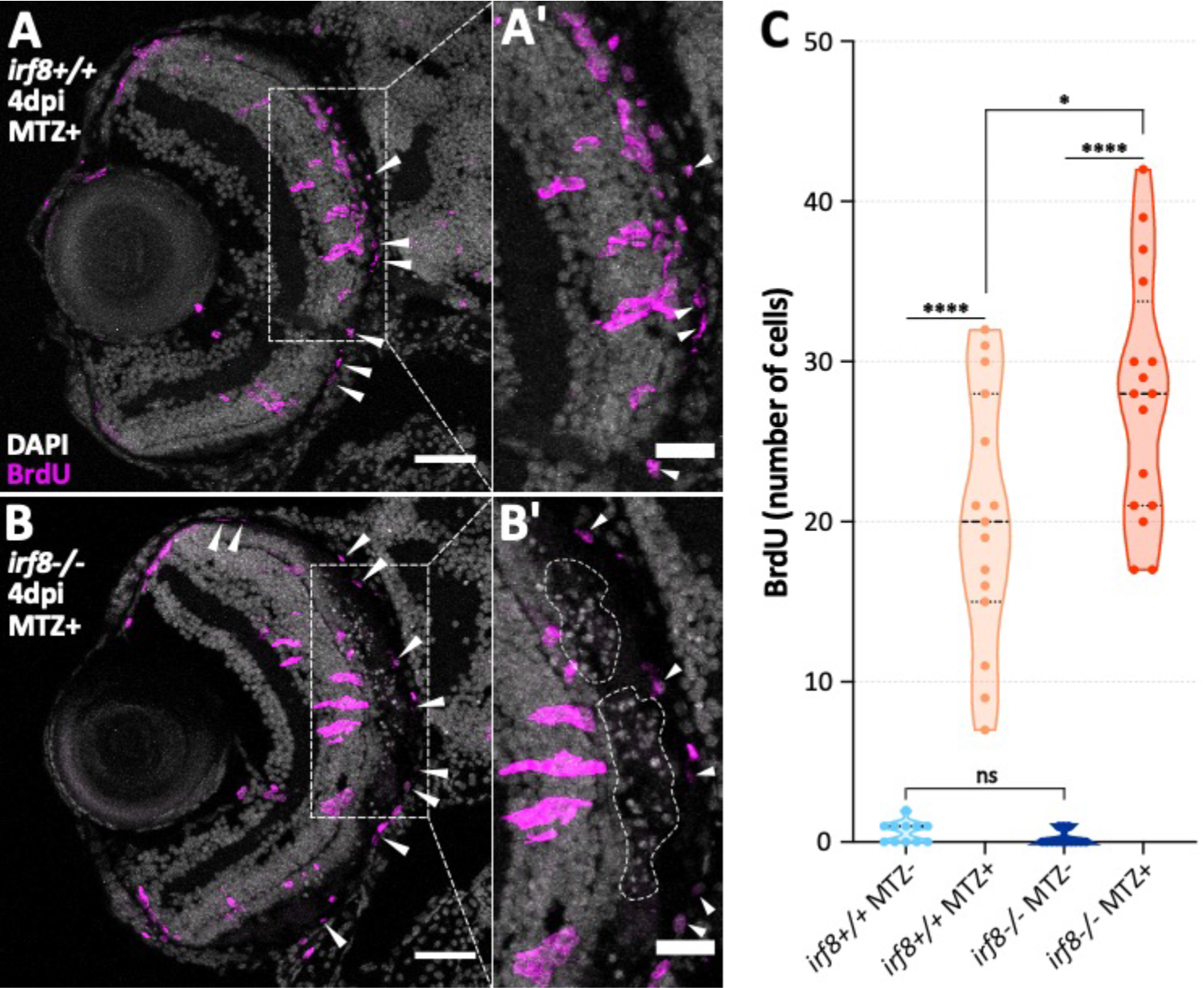
*irf8* mutant larvae retain proliferating cells and pyknotic nuclei post-RPE ablation. Fluorescent confocal micrographs of transverse sections from 4dpi MTZ+ ablated *irf8* wild-type **(A)** and *irf8* mutant **(B)** larval eyes. Images show peripherally localized BrdU^+^ cells in the *irf8* mutant larvae. Digital zooms highlight clusters of pyknotic nuclei retained between the outer nuclear layer and the RPE in *irf8* mutants (**B**’; *white dashed lines*) but not in *irf8* wildtype control siblings **(A’)**. *White arrowheads* mark BrdU-labeled cells (*magenta*) within the RPE layer. *White* (DAPI) labels nuclei. Scale bars represent 40μm **(A,B)** or 20μm **(A’,B’)**. **(C)** Violin plots showing quantification of the number of BrdU^+^ cells in the RPE layer in 9dpf MTZ-unablated and 4dpi MTZ+ ablated larvae. Data reveal a significant increase in proliferative cells in MTZ+ ablated *irf8* mutants when compared to MTZ+ ablated wild-type siblings (p=0.0111). Significantly more proliferative cells were present in both MTZ+ ablated *irf8* wild-types and mutants when compared to respective MTZ-unablated controls (p≤0.0001 for both comparisons). There was no significant difference between MTZ-unablated *irf8* wild-type and mutant larvae (p=0.1054). *Dashed black lines* represent the median and *dotted black lines* represent quartiles. Statistics (number of experiments, biological replicates (n), statistical test, and p-values) can be found in Table 2. Dorsal is up; definitions as follows: *, p-value ≤0.05; ****, p-value ≤0.0001; BrdU, bromodeoxyuridine; dpf, days post-fertilization; dpi, days post-injury; MTZ, metronidazole; ns, not significant; RPE, retinal pigment epithelium.

To look more closely at cell death in *irf8* mutants, we utilized terminal deoxynucleotidyl transferase dUTP nick end labeling (TUNEL) to identify cells undergoing programmed cell death in ablated *irf8* wild-type and mutant larvae. At 4dpi, accumulation of TUNEL^+^ puncta was apparent between the outer plexiform layer (OPL) and the basal side of the RPE in *irf8* mutants (Fig. 7B,D) when compared to wild-type siblings (Fig. 7A,C). In *irf8* mutants, there was a range of TUNEL^+^ staining, and we have shown representative sections (Fig. 7A,B) alongside extreme cases of TUNEL accumulation (Fig. 7C,D) for both *irf8* wild-type and mutant larval pools; these specific datapoints are labeled in Fig. 7E. Quantification of the total number of TUNEL^+^ puncta revealed no significant differences between unablated and ablated *irf8* wild-type larvae, nor between unablated *irf8* wild-type and mutant siblings (Fig. 7E; p=0.1694 and p=0.3626, respectively). The latter of these findings is important as this shows that the retention of dying cells was a result of MTZ-dependent RPE ablation, not the mutation in *irf8* itself. There were significant differences in the number of TUNEL^+^ puncta between ablated *irf8* mutant and wild-type siblings, and between ablated and unablated *irf8* mutants (Fig. 7E; p=0.001 and p<0.0001, respectively), indicating that dying cells are specifically retained in the damaged RPE of *irf8* mutants, supporting a role for MΦ/μglia cells in their removal.

**Figure 7.**
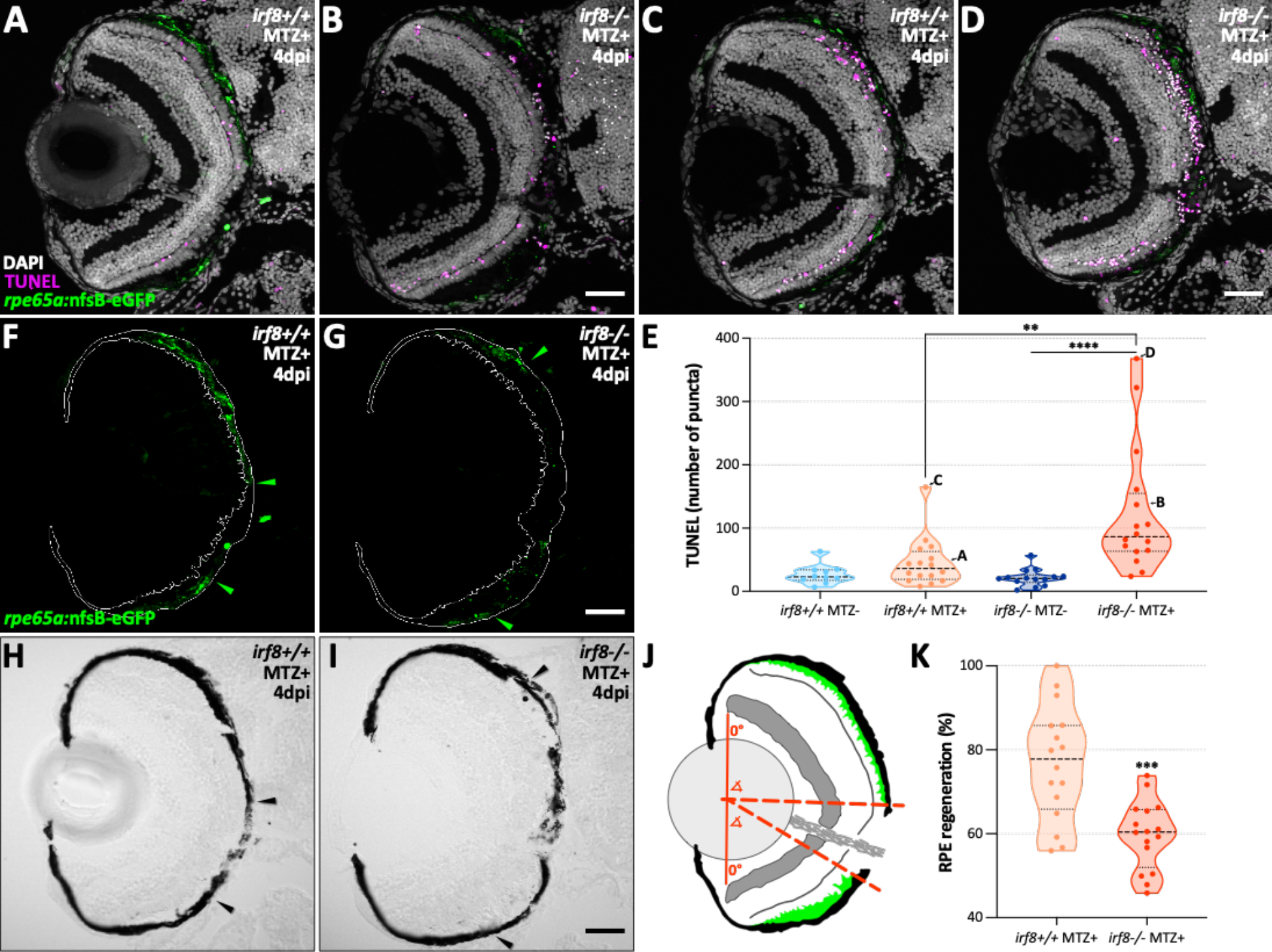
*irf8* mutant larvae show impaired RPE regeneration. **(A-D)** Fluorescent confocal micrographs showing TUNEL staining (*magenta*) on transverse sections from 4dpi MTZ+ ablated *irf8* wild-type **(A,C)** and *irf8* mutant **(B,D)** larval eyes. Images depict representative larvae **(A,B)** as well as larvae with the most TUNEL^+^ puncta from MTZ+ ablated datasets **(C,D)**. In both representative and extreme cases, images show more TUNEL^+^ puncta in the *irf8* mutants. *White* (DAPI) labels nuclei and *green* labels endogenous *rpe65a*:nfsB-eGFP. **(E)** Violin plots showing quantification of the number of TUNEL^+^ puncta located between the outer plexiform layer and the RPE in 9dpf MTZ-unablated and 4dpi MTZ+ ablated larvae. Data reveal a significant increase in TUNEL labeling in MTZ+ ablated *irf8* mutants when compared to MTZ+ ablated wild-type and MTZ-unablated mutant siblings (p=0.001 and p≤0.0001, respectively). There was no significant difference between MTZ-unablated and MTZ+ ablated *irf8* wild-type siblings or between MTZ- unablated *irf8* wild-type and mutant siblings (p=0.1694 and p=0.3626, respectively). **(F,G)** Fluorescent confocal micrographs of representative larvae **(A,B)**, showing area of the RPE (*white line*) and endogenous *rpe65a*:nfsB-eGFP expression (*green). Green arrowheads* delineate where peripheral-to-central continuous eGFP expression ends. **(H,I)** Brightfield confocal micrographs showing pigmentation relative to borders of continuous eGFP expression (*black arrowheads*). Images show impaired central recovery of intact eGFP and pigment in *irf8* mutants. All scale bars represent 40μm. **(J)** Schematic showing how RPE regeneration measurements were quantified. A dorsal-ventral line with midpoint (*solid red line*) was drawn and dorsal and ventral angle measurements were made between 0° and the point where continuous eGFP expression stopped (*dashed red lines*). **(K)** Violin plots showing a significant decrease in RPE regeneration in MTZ+ ablated *irf8* mutants when compared to MTZ+ ablated wild-type siblings (p=0.002). In all violin plots, *dashed black lines* represent the median and *dotted black lines* represent quartiles. Statistics (number of experiments, biological replicates (n), statistical test, and p-values) can be found in Table 2. Dorsal is up; definitions as follows: **, p-value ≤0.01; ***, p-value ≤0.001; ****, p-value ≤0.0001; dpf, days post-fertilization; dpi, days post-injury; MTZ, metronidazole; RPE, retinal pigment epithelium; TUNEL, terminal deoxynucleotidyl transferase dUTP nick end labeling.

To quantify RPE regeneration, we utilized the recovery of *rpe65a*:nfsB-eGFP transgene expression as a marker of RPE (25). RPE-ablated *irf8* wild-type larvae displayed a more centralized recovery of continuous endogenous eGFP expression when compared to RPE-ablated *irf8* mutants (Fig. 7F,G; *green arrowheads*). When the edges of eGFP recovery were overlaid onto brightfield images, lack of RPE tissue integrity in the central injury site (between the *black arrowheads*) was visible and the injury site appeared larger in *irf8* mutants (Fig. 7H,I). To quantify RPE regeneration, dorsal and ventral angle measurements were made based on the edges of continuous eGFP expression in each larva (Fig. 7J). Results showed that RPE regeneration was significantly decreased in ablated *irf8* mutants when compared to wild-type sibling controls (Fig. 7K; p=0.0002). Collectively, these results indicate that MΦs/μglia are required for the timely progression of RPE regeneration.

## DISCUSSION

Harnessing the intrinsic ability of the RPE to self-repair is an attractive alternative or supplemental therapeutic approach to mitigating RPE degenerative diseases, such as STGD, RP, and AMD. Little is currently known about the signals driving RPE regeneration due to the difficulty of studying intrinsic RPE repair in mammals, which are limited in regenerative capacity, and, until recently, the absence of a useful vertebrate model system in which intrinsic RPE regeneration could be studied. Recent characterization of a zebrafish RPE injury model has enabled us to begin to understand the molecular pathways regulating intrinsic RPE regeneration in a vertebrate system (25). Here, we provide strong evidence of immune system involvement during RPE regeneration. Specifically, we found that immune-related genes are upregulated in regenerating RPE, that MΦs/μglia are the responsive leukocyte post-RPE injury, and that inflammation and MΦ/μglia cells, specifically, are required for RPE regeneration.

As almost nothing is known about the molecular underpinnings of RPE regeneration, our initial approach was to perform an RNA-seq screen on FACS-isolated RPE to get a global picture of the changing gene expression profiles taking place after tissue damage. Of interest immediately was the observed upregulation of known recruitment factors for neutrophils (e.g. *cxcl8* and *cxcl18*) and macrophages (e.g. *il34*) in RPE at early- and peak-regenerative timepoints (e.g. 2dpi and 4dpi). While robust neutrophil recruitment was not observed, there was significant infiltration of MΦs/μglia from 2-4dpi. Additionally, transcriptome profiling of MΦs/μglia revealed a significant upregulation of cell cycle-related genes at 2 and 4 days post-RPE ablation. As IL-34 has been shown to promote survival and proliferation in macrophages and microglia (48), it is plausible that damaged RPE express *il34* to recruit MΦs/μglia for tissue repair. In support of this assertion, O’Koren *et al*. recently identified two microglia niches in the mouse retina that vary by dependence on IL-34 under homeostatic conditions (33). After photoreceptor damage, a majority of microglia from both niches migrated to the apical side of the RPE; however, it was the IL-34-dependent microglia that functioned to protect the RPE and maintain the integrity of the blood-retinal-barrier (33). The photoreceptors and RPE exist as a functional unit, and damage to one tissue can elicit an injury response in the other (1). While O’Koren *et al*. did not determine if RPE was a source of IL-34 post-photoreceptor damage in the mouse, the fact that IL-34-dependent microglia home to RPE under stress suggests this is a possibility in mammals.

*il11b* was the most significantly upregulated DEG in FACS-isolated RPE at 2dpi. Interleukin 11 (IL-11) expression is induced by inflammation and has been shown to have a range of functions, including as an anti-inflammatory cytokine and a survival and differentiation factor in the CNS (70). IL-11 also drives proliferation of Müller glia-derived progenitor cells (MGPCs), which proliferate after retinal injury to regenerate lost neurons in teleost fish (15–17). In the zebrafish retina, both *il11a* and *il11b*, along with their receptor, *gp130*, are expressed in MGPCs, and recombinant mammalian IL-11 was shown to promote a modest increase in MGPC proliferation and reprogramming in the absence of retinal injury (71). Recombinant IL-11 was more effective in combination with active leptin signaling (71), indicating an added level of control post-injury, perhaps to induce proliferation only in specific cell types (e.g. MGPCs). Interestingly, our RPE RNA-seq results showed that leptin (*lepb*) is upregulated alongside *il11b* at 2dpi (Table S1). While the source of newly regenerated RPE in our model remains unknown, previous characterization has shown that it is likely residual peripheral RPE that respond by proliferating post-injury (25). Evidence for an RPE stem cell exists from studies in medaka (72) and in adult human donor RPE, where some cells could be induced to proliferate *in vitro* (22), but whether the zebrafish proliferating peripheral RPE are stem cell-like is not yet known. Nevertheless, expression of *il11b* and *lepb* by early-stage regenerating RPE is compelling and requires further examination.

We also noted significant upregulation of MMPs post-RPE injury in both RPE and MΦ/μglia RNA-seq datasets. Specifically, in RPE, *mmp13b* was upregulated at 2dpi and *mmp9/13b* at 4dpi (Tables S1 and S2); in MΦs/μglia, *mmp9/13a* was upregulated at 2dpi (Table S7). MMP9/13 are of interest in the context of regeneration, as expression has been reported in a range of tissues undergoing repair after injury, including: limb (73), muscle (74), liver (75), heart (28), fin (76–78), and retina (67, 79). In all of these systems, which span mammalian and non-mammalian vertebrates, disruption of MMP9 and/or MMP13 impeded the normal progression of tissue repair, highlighting the importance of these proteases in a range of regenerative contexts. Our findings suggest that the RPE may also utilize MMPs to mediate regeneration.

Leukocytes of the innate immune response (e.g. neutrophils and macrophages) predominate in zebrafish at the larval ages used for this study. In a typical response to inflammation, tissue resident macrophages precede neutrophils at the injury site, which are followed by recruited macrophages (80). Neutrophils play an important role in pathogen containment and mitigating infection, while macrophages are critical for tissue repair (80). Here, we showed no appreciable accumulation of neutrophils at any of the timepoints queried; instead, MΦs/μglia were the dominant leukocyte located in the RPE postablation. These results align with regenerative studies in the retina employing targeted genetic ablation (35, 81) or inducing damage with a cytotoxic agent (36); while regenerative studies using needle poke in the eye (35), and in non-ocular tissues utilizing other approaches (e.g. tissue resection or cryoinjury), have showed robust infiltration of neutrophils followed by macrophage influx (27–29, 82, 83). The range in leukocyte responses to damage across injury paradigms may be based on the extent of the injury (e.g. resection of whole tissue versus genetic ablation of a single cell type), the type of tissue, and the location within the organism (i.e. risk of pathogen exposure); likely all of these are contributing factors. Indeed, studies where the injury site is exposed to the external environment (e.g. caudal tail fin resection) reported robust neutrophil infiltration (29, 82, 83). Interestingly, impeding neutrophil infiltration had little effect on tissue regeneration (29, 82), indicating these cells may not be directly involved in tissue repair, consistent with our observation that neutrophils do not readily accumulate in the RPE post-ablation.

Macrophages and microglia are extremely dynamic cells that adjust to fit their microenvironment. In the CNS, microglia under homeostatic conditions extend cellular processes into the surrounding tissue and take on a ramified morphology; in response to injury or inflammation, these same cells rapidly retract processes and round up, appearing more amoeboid in morphology (84). Likewise, changes in morphology are often related to functional roles, such as phagocytosis (56). In addition to increased infiltration post-RPE ablation, we also qualitatively observed distinct morphology changes in 4C4^+^ MΦs/μglia in whole mount tissue. Specifically, we saw a more amoeboid phenotype in 4C4^+^ cells at 2dpi, 3dpi, and 4dpi when compared to unablated controls, which showed a more ramified morphology and significantly fewer MΦs/μglia localized to the RPE. This observation aligns with previous findings that microglia round up in response to damage in zebrafish (35, 36, 81) as well as in mammalian (85) CNS tissues. Further, our RNA-seq results indicate that, post-RPE ablation, MΦs/μglia express markers of phagocytosis (e.g. *anxa1a* (61, 62)) and cell cycle-related genes, the latter of which has been shown to be a signature of homeostatic amoeboid microglia in rat CNS tissue (86). Quantitatively, we detected a significant increase in the sphericity of anterior MΦs/μglia during peak infiltration at 3dpi. We chose to measure sphericity in the entire population of anterior MΦs/μglia as previous studies have reported reactivity of phagocytic cells outside the immediate injury site (35, 81). It is possible that more localized sphericity differences exist at 2dpi and 4dpi that are diluted by the entire population of anterior MΦs/μglia; however, more stringent characterization of cells at higher magnification *in vivo* will be required to answer this question.

Several phases have been characterized following tissue injury: I) the inflammatory phase, in which leukocytes are recruited, secrete pro-inflammatory cytokines, and begin phagocytosis; II) the resolution phase, in which macrophages continue to phagocytose cell debris and make the switch from a pro- to an anti-inflammatory phenotype; and III) the regeneration phase, in which injured tissues initiate proliferation (82, 87). Indeed, sequential progression of resolution and regeneration phases have been shown post-tail fin resection in zebrafish (82). We propose that similar phases exist during RPE regeneration (summarized in Fig. 8). Our previous characterization of RPE regeneration in zebrafish showed that, between 3-4dpi, proliferation peaked in the RPE layer and pigmentation was recovered (25). We showed evidence here that MΦs/μglia begin to infiltrate the RPE and may be phagocytic by 2dpi and observed peak infiltration of MΦs/μglia in the RPE injury site at 3dpi, which waned by 4dpi; representing a potential window when inflammation is resolved (Fig. 8; 2-4dpi). Building off of these data, it appears that functional MΦ/μglia presence in the injury site precedes as well as overlaps with peak RPE proliferation and visible recovery of pigmentation; thus, 3-4dpi may represent a critical window post-RPE ablation when the resolution phase ends and the regeneration phase begins (Fig. 8).

**Figure 8.**
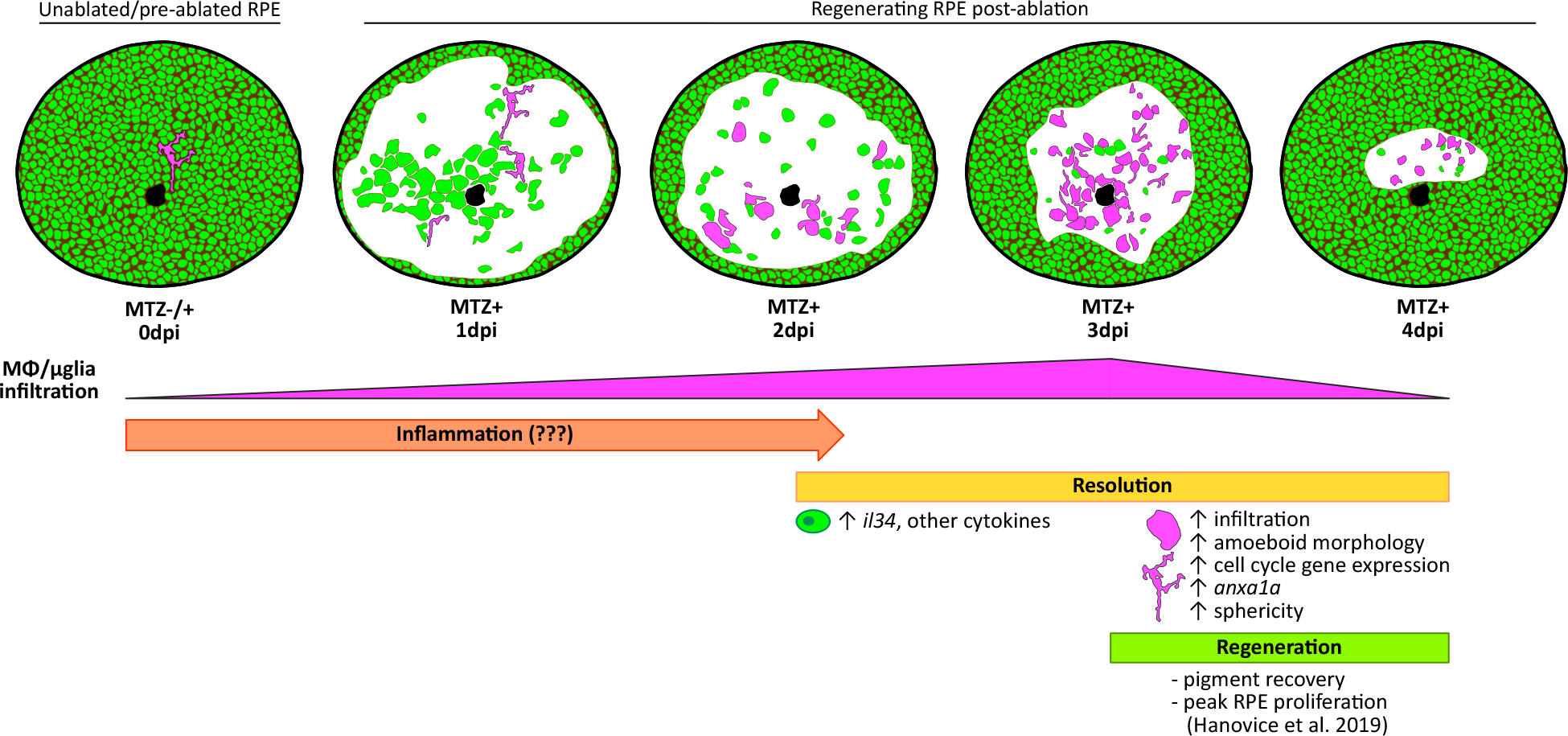
Phases of immune involvement during RPE regeneration. Schematic showing few ramified MΦs/μglia (*magenta*) present in the RPE (*green*) of unablated larvae. Infiltration of MΦs/μglia to the central RPE injury site post-ablation begins at 2dpi, peaks at 3dpi, and wanes by 4dpi, representing a window of time when inflammation is likely resolved (2-4dpi; *yellow*). During resolution, MΦs/μglia appear amoeboid in morphology, express cell cycle and phagocytosis markers (e.g. *anxa1a*), and RPE express *il34* and other cytokines. Previous characterization has shown peak proliferation in the RPE layer and recovery of pigment between 3-4dpi (25). This coupled with the decreased presence of MΦs/μglia in the RPE by 4dpi may hint to a window of time post-ablation (e.g. 3-4dpi; *green*) when inflammation has been resolved, enabling peak RPE regeneration. Definitions as follows: dpf, days postfertilization; dpi, days post-injury; MΦs/μglia, macrophages/microglia; MTZ, metronidazole; RPE, retinal pigment epithelium.

In agreement with previous reports in multiple reparative contexts (e.g. (27, 29, 35, 67, 82)), we demonstrate that inflammation and MΦs/μglia are required for RPE regeneration *in vivo*. Synthetic GCs, such as dexamethasone, have been widely used to suppress inflammation and do so by attenuating the inflammatory phase post-injury and driving macrophages toward an anti-inflammatory phenotype (87–89). Results from our study support the existence of a critical inflammatory phase during RPE regeneration, as evidenced by expression of phagocytic (e.g. *anxa1a*) and pro-inflammatory genes (e.g. *cxcl8a, cxcl18b*) and recruitment of MΦs/μglia to the ablated RPE (Fig. 8). Inhibition of inflammation using dexamethasone resulted in decreased proliferation in the RPE layer and delayed recovery of a pigmented monolayer. These findings align with studies in the zebrafish retina, which showed less proliferative MGPCs and reduced photoreceptor regeneration post-dexamethasone treatment (35, 67).

Using an *irf8* mutant, which lacks macrophages until around 7dpf and microglia until at least 31dpf (40), we showed that RPE regeneration is impaired post-ablation. Retention of pyknotic nuclei and TUNEL^+^ debris in and adjacent to the RPE injury site was a prominent phenotype in RPE-ablated *irf8* mutants. Previously, we have shown significant cell death in response to MTZ-mediated RPE ablation in both the RPE and the photoreceptors as early as 12 hours post-injury (hpi) and 18hpi, respectively (25); and it is likely that debris remains in *irf8* mutants due to the lack of effective clearance by MΦs/μglia, similar to phenotypes in *irf8* mutants post-spinal cord injury (27). MΦ/μglia infiltration was unaffected and pyknotic nuclei did not accumulate in dexamethasone-treated larvae, indicating intact phagocytic function, yet the outcome was still impaired RPE regeneration. GCs have been shown to have varying effects on macrophage migration (89); indeed, in GC-treated zebrafish post-wounding, leukocyte migration defects have been detected in some injury paradigms (27, 35, 67), but not others (90, 91). Further, GCs have been shown to increase phagocytosis by macrophages to promote expedited resolution of inflammation (87, 89), which may explain the lack of pyknotic nuclei accumulation in dexamethasone-treated larvae post-RPE damage. Collectively, the results from dexamethasone and *irf8* mutant experiments hint at the existence of critical inflammatory and resolution phases post-RPE ablation, and we provide evidence that by-passing either phase results in the same outcome: impairment of RPE regeneration.

Lastly and paradoxically, we observed an increase in proliferating cells in the RPE of *irf8* mutants post-ablation. While increased proliferation may seem beneficial in the context of regeneration, RPE overproliferation can lead to pathology (e.g. in proliferative vitreoretinopathy (19)). We have shown that proliferation tapers off in the RPE at 4-5dpi (25), so prolonged proliferation is likely not advantageous for RPE regeneration. In support of this, despite the presence of more BrdU^+^ cells, RPE regeneration is impaired in *irf8* mutants. A possible explanation for the increase in proliferation is the lingering TUNEL^+^ cells, as signaling via apoptotic bodies has been shown to induce proliferation in different injury contexts (92, 93). The emergence of apoptotic and proliferating cells coincides within the first 24-hours post-RPE injury (25), thus it is possible that apoptosis is an additional signaling factor driving proliferation. This assertion, while remaining to be tested, is an interesting avenue for future study.

## MATERIALS & METHODS

### Zebrafish lines, maintenance, and husbandry

All zebrafish embryos, larvae, and adults were handled in compliance with the regulations enforced by the University of Pittsburgh Institutional Animal Care and Use Committee (IACUC). All zebrafish lines used for this study are listed in Table 1. Lines harboring fluorescent transgenes were screened using an AxioZoom 16 Fluorescent Stereomicroscope (Zeiss, Oberkochen, Germany). Zebrafish used for line maintenance and spawning were housed in a facility kept at 28.5°C on a 14-hour light/10-hour dark cycle. Embryos collected for experimentation were housed in an incubator kept at 28.5°C (no light/dark cycle) until euthanasia by tricaine (MS-222; Fisher Scientific, Pittsburgh, Pennsylvania) overdose. Larvae euthanized on or before 9 days post-fertilization (dpf) were unfed (Hernandez et al., 2018), while those aged 9dpf or older were fed a Zeigler AP100 size 2 powder diet (Zeigler Bros, Inc., Gardners, Pennsylvania). Zebrafish adults and larvae from *st95 (irf8* mutation) crosses were genotyped as described using published primers (Table 1; (40)). For genotyping, fresh tissue in 50mM NaOH was heated to 95°C for 10 minutes. Once cooled to 4°C, Tris buffer (pH 8, 100mM final concentration) was added for storage at −20°C. Genotyping PCR products were digested with AvaI restriction enzyme (Abcam, Cambridge, Massachusetts) for 1 hour at 37°C and electrophoresed on a 3% agarose gel in 1X Tris-acetate-EDTA (TAE) buffer.

**Table 1.**
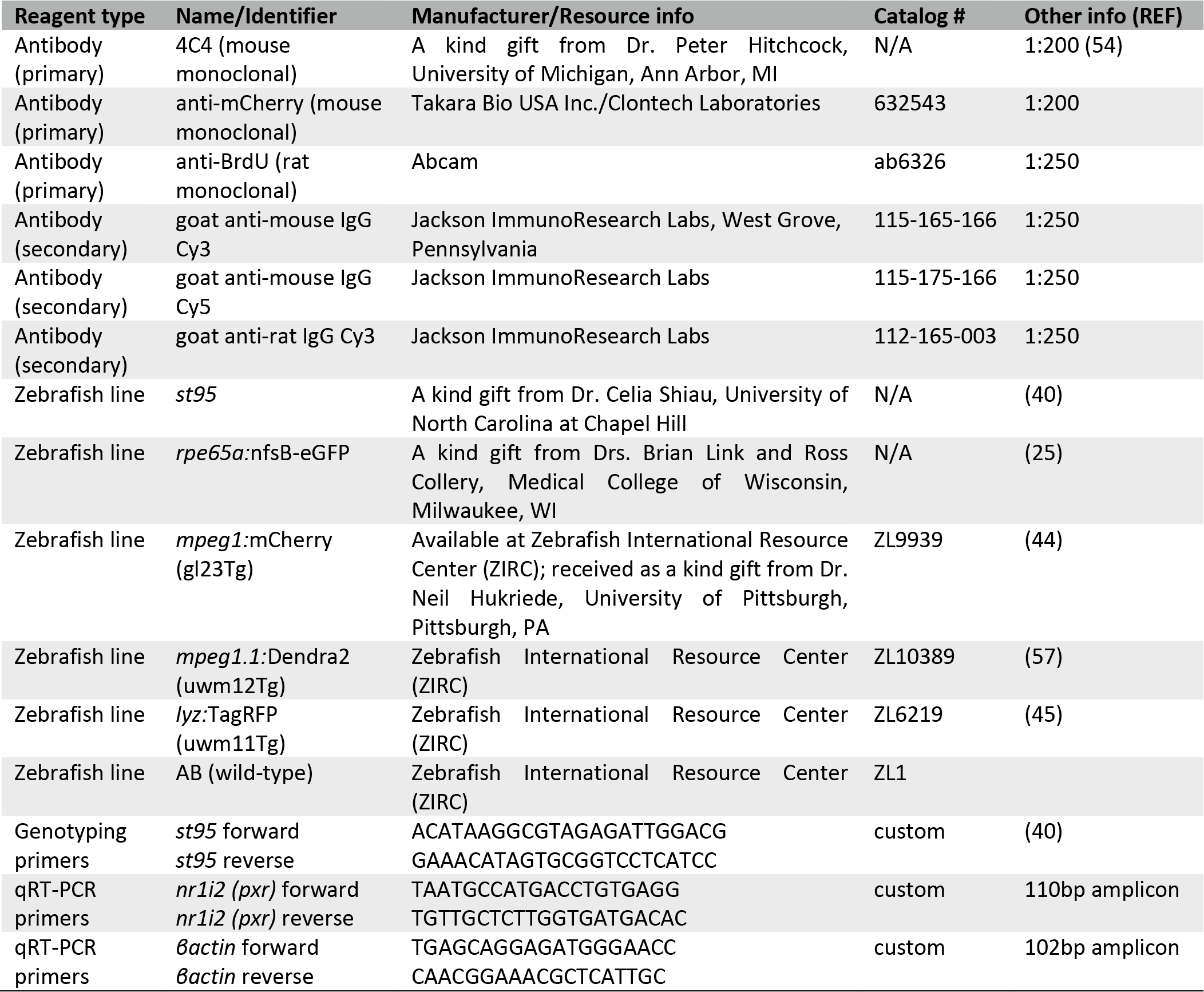
Resources and reagents used.

### Zebrafish retinal pigment epithelium (RPE) ablation paradigm

RPE ablation was performed as described previously (25). Briefly, embryos collected from *rpe65a*:nfsB-eGFP outcrosses were incubated in system water treated with 1.5X n-phenylthiourea (PTU; Sigma-Aldrich, St. Louis, Missouri) from 0.25 – 5 days post-fertilization (dpf) to prevent melanogenesis. On 5dpf, larvae were tricaine-anesthetized, screened for eGFP, and subsequently treated with 10mM metronidazole (MTZ; Sigma-Aldrich) in system water for 24 hours to ablate the RPE. The MTZ solution was made on 5dpf, immediately prior to screening, by dissolution in system water at 37°C (shaking) for 1 hour, with a subsequent “cooling” step at room temperature for 1 hour. MTZ was washed out after 24 hours (designated 1dpi) and larvae remained in system water until fixation.

### Dexamethasone treatment

The timing of dexamethasone treatment closely follows that of White *et al*. (35). On 4dpf, larvae were incubated in system water with 1.5XPTU treated with 50μM dexamethasone (Sigma-Aldrich) in dimethylsulfoxide (DMSO) or 0.05% DMSO (vehicle control) for 24 hours (pre-ablation treatment). The next day (5dpf), larvae were ablated as described above in system water with 10mM MTZ treated with 50μM dexamethasone or 0.05% DMSO. After 24 hours (1dpi), MTZ was washed out and larvae remained in system water treated with 50μM dexamethasone or 0.05% DMSO until fixation. Water changes were performed daily to replenish dexamethasone and DMSO.

### Single cell dissociation and fluorescence activated cell sorting (FACS) for RNA-sequencing

For both methods described below (modified from (94, 95)), zebrafish larvae were euthanized and transferred to ice cold 1XPBS for bilateral enucleations. Enucleations were performed using chemically-sharpened tungsten needles and larval eyes were pooled for each experiment. Additionally, for both methods, gated cells were sorted directly into reagents for cDNA synthesis (see bulk RNA-sequencing methods) in a 96-well plate. All sorting was performed by the Flow Cytometry Core at the University of Pittsburgh School of Medicine Department of Pediatrics.

#### RPE cell isolation

Experiments used embryos collected from *rpe65a*:nfsB-eGFP x wild-type AB outcrosses and were performed in biological triplicate on unablated (MTZ-) and ablated (MTZ+) siblings at: 7dpf (n=55, n=25, n=25); 2dpi (n=55, n=25, n=25); 9dpf (n=55, n=37, n=35); 4dpi (n=55, n=37, n=38); 12dpf (n=45, n=50, n=35); and 7dpi (n=35, n=40, n=28). Pooled eyes were incubated in 1.5g/L collagenase for 50-60 minutes at 28.5°C with occasional, gentle trituration and observation using a stereomicroscope. When the RPE was visibly sloughing off, cell suspensions were pelleted at 450 x g for 5 minutes. Cells were resuspended in trypsin-EDTA (0.25%) and incubated for 10-15 minutes at 28.5°C, as described for collagenase. The resulting cell suspension was pipetted through a 70μm strainer into ice cold 1XPBS with 2mM CaCl2 and 5% fetal bovine serum (FBS; to stop the trypsin reaction) and pelleted at 450 x g for 5 minutes at 4°C. Cell pellets were washed with ice cold 1XPBS 3 times (spinning at 450 x g for 5 minutes at 4°C between washes). During the second wash, cell pellets were resuspended in ice cold 1XPBS with diluted LIVE/DEAD Fixable Aqua Dead Cell Stain (1:1000; Invitrogen, Waltham, Massachusetts) and incubated on ice for 30 minutes in the dark. Live (Aqua-dim), eGFP^+^ cells were isolated using a FACSAria IIu cell sorter (BD Biosciences, Franklin Lakes, New Jersey). A single gate was set by eGFP intensity in unstained unablated (MTZ-) and ablated (MTZ+) *rpe65a*:nfsB-eGFP^+^ samples and used to sort all biological triplicates.

#### Macrophage/microglia cell isolation

Experiments used embryos collected from *rpe65a*:nfsB-eGFP x *mpeg1*:mCherry outcrosses and were performed in biological triplicate on unablated (MTZ-) and ablated (MTZ+) siblings at: 7dpf (n=40, n=38, n=50); 2dpi (n=35, n=36, n=35); 9dpf (n=32, n=35, n=36); and 4dpi (n=24, n=30, n=41). To minimize exposure of MΦs/μglia to prolonged enzyme digestion prior to sorting, some steps used for the RPE dissociation protocol above were modified. Briefly, pooled eyes were moved directly into trypsin-EDTA (0.25%) and mechanically dissociated using a 27 ½ gauge syringe needle. Cell straining, washing, and LIVE/DEAD labeling steps were performed as described above for RPE. Live (Aqua-dim), mCherry^+^ cells were isolated using a FACSAria IIu cell sorter (BD Biosciences). A single gate was initially set by mCherry intensity in unstained and *mpeg1*:mCherry^+^ (*rpe65a*:nfsB-eGFP^-^) samples and then finalized using unablated (MTZ-) and ablated (MTZ+) *mpeg1*:mCherry^-^ (*rpe65a*:nfsB-eGFP^+^) samples to prevent collection of autofluorescent cells; this gate was used to sort all biological triplicates.

### Bulk RNA-sequencing and bioinformatics

#### Library preparation and sequencing

All library preparation, quality control analyses, and next generation sequencing was performed by the Health Sciences Sequencing Core at Children’s Hospital of Pittsburgh. Synthesis, amplification, and purification of cDNA was performed using either the SMART-Seq v4 Ultra Low Input RNA Kit (Takara Bio USA, Inc., Mountain View, California) for RPE samples or the SMART-Seq HT Ultra Low Input RNA Kit (Takara Bio USA, Inc.) for MΦ/μglia samples, both of which require ≤1000 cells to generate sequencing-quality cDNA. The resulting double-stranded cDNA was validated using an Advanced Analytical Fragment Analyzer 5300 or TapeStation 2200 system (both Agilent Technologies, Inc., Santa Clara, California) and quantified using a Qubit 4 Fluorometer (Invitrogen). High-sensitivity reagents were used to validate and quantify cDNA and only high-quality samples were submitted for library preparation and sequencing. cDNA sequencing libraries were prepared using the Nextera XT DNA Library Preparation Kit (Illumina, Inc., San Diego, California) following manufacturer instructions, and 75 cycle, 2×75bp paired-end read sequencing runs were performed on a NextSeq 500 system (Illumina, Inc.) using High-Output 150 flow cells, aiming for 40 million reads per sample.

#### Bioinformatics analysis

Fastq files containing raw read data were obtained from the Health Sciences Sequencing Core and imported into CLC Genomics Workbench (Qiagen Digital Insights, Hilden, Germany) licensed through the Molecular Biology Information Service of the Health Sciences Library System at the University of Pittsburgh. CLC Genomics Workbench was used for all subsequent read processing, mapping, and differential expression analyses. Briefly, imported Illumina reads were trimmed of adapters and subjected to quality control analysis. Read quality assessment was based on Phred-scores (96), and all samples showed the majority of read sequences averaging Phred-scores ≥20. Trimmed reads were mapped to *Danio rerio* reference genome assemblies (GRCz10 for the RPE dataset and GRCz11 for the MΦ/μglia dataset) and mapped gene tracks were analyzed for differential gene expression (DEG) with unablated (MTZ-) alignments set as the control group.

#### Pathway enrichment analysis

Filtering was performed on DEGs from 7dpf/2dpi and 9dpf/4dpi RPE and MΦ/μglia datasets; only genes with maximum group mean ≥5, fold change ≥2, and false discovery rate (FDR) p-value ≤0.05 were submitted for pathway enrichment analysis using the PANTHER statistical overrepresentation test (http://www.pantherdb.org/; released April 7, 2020). Lists of filtered genes were uploaded with *Danio rerio* set as the reference genome and Reactome pathways as the annotation source (version 65, released December 22, 2019). Results for RPE represent 332 mapped genes from the 7dpf/2dpi dataset and 301 mapped genes from the 9dpf/4dpi dataset; results for MΦs/μglia represent 67 mapped genes from the 7dpf/2dpi dataset and 208 mapped genes from the 9dpf/4dpi dataset. The Fisher’s Exact test was performed with FDR correction for each analysis and only enriched pathways with FDR p-value <0.05 and >5 genes/pathway were examined. Heatmaps were generated in CLC Genomics Workbench using filtered gene tracks based on enriched pathway gene sets.

#### Data Availability

Raw (fastq) and processed (total counts and transcripts per million (TPM) values) RNA-seq data will be accessible in NCBI Gene Expression Omnibus (GEO) (97, 98) upon publication.

### Quantitative real-time PCR (qRT-PCR)

Whole 5dpf zebrafish larvae treated with 50μM dexamethasone (3 separate experiments: n=24, n=31, n=19) or 0.05% DMSO (3 separate experiments: n=23, n=30, n=18) for 24 hours were euthanized and pooled for each experiment. Pooled tissue was homogenized in Buffer RLT Plus by vortexing and total RNA was purified using the RNeasy Plus Mini Kit (Qiagen). cDNA was generated using the iScript cDNA Synthesis Kit (Bio-Rad Laboratories, Hercules, California) and qRT-PCR reactions were run on a CFX384 Touch Real-Time PCR Detection System (Bio-Rad Laboratories) using the iTaq Universal SYBR Green Supermix (Bio-Rad Laboratories) and primers designed to amplify 110bp of the *nr1i2* (*pxr*) coding region spanning exons 2-3 (Table 1; NCBI reference sequence: NM_001098617.2). Pooled experiments were performed in biological triplicate and C_T_ values from each experiment represent the average of 3 technical replicates to account for well-to-well variation. Fold change was calculated using the comparative C_T_ method (2^−ΔΔCT^; (99)) and *βactin* was used as the reference gene. *βactin* C_T_ values did not show evidence of treatment-dependent differences (100).

### Immunohistochemistry

For both methods described below, zebrafish larvae were euthanized and fixed in 4% paraformaldehyde (PFA) either at room temperature for 2-3 hours or at 4°C overnight.

#### Sectioned tissue

Post-fixation, larvae were prepared for cryosectioning and stained using a previously described method, with modification (101). Larvae were washed once with 1XPBS, sucrose protected using a 25% to 35% gradient, and embedded in optimal cutting temperature (OCT) compound prior to cryosectioning. Transverse sections were acquired at 12μm thickness on a Leica CM1850 cryostat (Leica Biosystems, Wetzlar, Germany) on poly-L-lysine-coated slides. Sections were rehydrated with 1XPBS for 5 minutes and subsequently blocked using 5% normal goat serum (NGS, heat-inactivated) in 1XPBS with 0.1% Tween-20 and 1% DMSO (PBTD) for at least 2 hours at room temperature. For BrdU staining, sections were rehydrated and underwent an antigen retrieval step (4N HCl incubation for 8 minutes at 37°C), then 3x 10-minute washes with PBTD prior to blocking. Primary antibody (see Table 1 for all antibodies and dilutions used for this study) was diluted in 5% NGS blocking solution and incubated at 4°C overnight. The next day, slides were washed for 10 minutes, 3 times with PBTD and secondary antibody diluted in 5% NGS blocking solution was added for 3 hours at room temperature. Slides were again washed for 10 minutes, 3 times with PBTD and counterstaining with DAPI in 5% NGS blocking solution took place after the first wash. Coverslips (No. 1) were mounted using Vectashield (Vector Laboratories, Burlingame, California) and sealed with nail polish. Slides were stored at 4°C until imaging.

#### Whole mount tissue

Post-fixation, larvae were stored in 1XPBS until staining, the methods for which were adapted from (102). Whole larvae were rinsed in 1XPBS with 0.1% Tween-20 (PBST), rinsed in deionized water, then permeabilized in 100% acetone at −20°C for 12 minutes and subsequently washed for 5 minutes in deionized water and rinsed once in PBST. To improve enucleations, larvae were enzyme-digested first in collagenase (1mg/mL in PBST) and then in proteinase K (2mg/mL in PBST); each incubation was for 30 minutes at room temperature. Post-enzyme digestion, larvae were re-fixed in 4% PFA for 20 minutes at room temperature, rinsed in PBST, rinsed in 1XPBS containing 1% bovine serum albumin (BSA), 1% DMSO, 0.5% Triton X-100, and pH-adjusted to 7.3 (PBDTX), and blocked using 2% NGS in PBDTX for at least 1 hour at room temperature. Primary antibody was diluted in 2% NGS blocking solution and incubated at 4°C overnight. The next day, larvae were washed (rotating) for 15 minutes, 4 times in PBDTX and secondary antibody diluted in 2% NGS blocking solution was added for 4 hours at room temperature. Larvae were again washed on a rotator for 15 minutes, 4 times in 1XPBS with 0.5% Triton X-100 (PBSTX), then rinsed with 1XPBS and counterstained with DAPI in 1XPBS. Larval eyes were enucleated using chemically-sharpened tungsten needles and mounted in 0.5% low-melt agarose between a slide and coverslip immediately prior to imaging. For larvae harboring the *lyz*:TagRFP transgene, background fluorescence was dampened by subjecting samples to all steps listed above with two modifications: I) 5% NGS replaced 2% NGS and II) endogenous TagRFP was imaged, thus no primary or secondary antibodies were used.

### Bromodeoxyuridine (BrdU) incorporation and terminal deoxynucleotidyl transferase dUTP nick end labeling (TUNEL) assays

For BrdU incorporation assays, zebrafish larvae were exposed to 10mM BrdU (Sigma-Aldrich) in system water for 24 hours prior to fixation at 9dpf/4dpi. TUNEL assays were performed on sectioned tissue using the TMR red In Situ Cell Death Detection Kit (Roche, Basel, Switzerland), following the manufacturer protocol for labeling of cryopreserved tissues. Tissue was counterstained with DAPI in 1XPBS immediately prior to mounting coverslips with Vectashield, as described above.

### Imaging data acquisition, processing, and quantification

#### Confocal microscopy

All confocal images were acquired using an Olympus Fluoview FV1200 laser scanning microscope (Olympus Corporation, Shinjuku, Tokyo, Japan). All larval sectioned tissue was imaged using a 40X (1.30 NA) oil immersion lens and all whole mount tissue was imaged using a 20X (0.85 NA) oil immersion lens (both lenses from Olympus Corporation); all z-stacks acquired had 1μm z-step intervals. Raw imaging data were quantified and processed for presentation in figures using FIJI (ImageJ; (103)). Cell and puncta count quantification (e.g. *lyz*:TagRFP, BrdU, TUNEL) was done manually using the Cell Counter plug-in in FIJI. Briefly, *lyz*:TagRFP^+^ cells were counted if overlapping with eGFP signal (RPE) in whole mounts; in sectioned tissue, BrdU^+^ cells within the RPE layer were counted. To quantify TUNEL^+^ puncta, background was subtracted (rolling ball radius, 50 pixels) from maximum-projected z-stacks and puncta between the distal-most edge of the outer plexiform layer and the basal side of the RPE were counted. Data from BrdU and TUNEL experiments represent the sum of cells or puncta from 3 consecutive central sections. Percent area quantification was performed on 8-bit maximum-projected z-stacks in FIJI and regions of interest (ROIs) were generated manually using the polygon selection tool. ROIs for whole mount images were designated based on pigment in the brightfield channel and lens background was subtracted from all images prior to measurement. ROIs for sectioned tissue were drawn to encompass the RPE monolayer, e.g. the space between the outer-most edge of the outer nuclear layer and the basal side of the RPE. Thresholding was performed only on the secondary antibody channel (e.g. 4C4, *mpeg1*:mCherry signals) and was set at 40/255 (8-bit) for all percent area quantification; images with high background were omitted from quantification. Data from *mpeg1*:mCherry experiments represent percent area measurements from the central-most section of each larva. Percent pigment recovery and percent RPE regeneration were quantified in similar ways utilizing FIJI software. Briefly, a dorsal-ventral-running reference line with midpoint was drawn from either the distal-most tip of BrdU^+^ signal in the ciliary marginal zones (CMZ; pigment recovery quantification) or the distal-most edge of the inner nuclear layers, adjacent to the CMZ (RPE regeneration quantification). Using the reference line and midpoint, dorsal and ventral angle measurements were made, summed, and converted to a percentage to determine the total amount of pigment recovery or RPE regeneration (e.g. 100% = 180°). Criteria for pigment recovery was based on the peripheral-to-central expansion of continuous (i.e. not patchy), darkly pigmented tissue and angle measurements were made at the junction where tissue became depigmented and visibly disorganized. Criteria for RPE regeneration was based on the peripheral-to-central expansion of continuous, eGFP^+^ tissue and angle measurements were made at the junction where eGFP signal stopped or became patchy or punctate. Data from pigment recovery and RPE regeneration experiments represent measurements from the central-most section of each larva.

#### In vivo light-sheet microscopy

Experiments were performed at the Marine Biological Laboratory in Woods Hole, MA and used embryos collected from *rpe65a*:nfsB-eGFP x *mpeg1.1*:Dendra2 outcrosses. All data were acquired using the Tilt light-sheet imaging system (Mizar Imaging, Woods Hole, Massachusetts; (104)) fitted to a Nikon Eclipse Ti2-U inverted microscope equipped with 488nm and 561nm lasers (Nikon Corporation, Shinagawa, Tokyo, Japan). The RPE of eGFP and Dendra2 double-positive larvae was ablated as described above. For imaging, larvae were tricaine-anesthetized and embedded within optically clear glass imaging cubes (Mizar Imaging) in embryo medium with 1% low melt agarose and 20% OptiPrep (Stemcell Technologies, Inc., Vancouver, British Colombia, Canada). Larvae were oriented anterior-facing the light sheet source and dorsal-facing the detection objective. Immediately prior to image acquisition, Dendra2 (green) was photoconverted to Dendra2 (red) by 1-minute exposure to an Intensilight C-HGFI (Nikon Corporation) ultraviolet laser. Successful conversion was confirmed and ~300μm z-stacks (1μm z-step interval) for each larva were acquired using a 20X (0.95 NA) water immersion lens and NIS-Elements software (both Nikon Corporation). 3D deconvolution was performed in NIS-Elements and deconvolved z-stack files were 3D rendered for sphericity quantification using Imaris version 9.5 (Bitplane, Belfast, United Kingdom). In Imaris, isosurfacing was performed with 561nm set as the source channel (*mpeg1.1*:Dendra2 (red)). A Gaussian filter (smoothing = 0.5μm) and absolute intensity threshold of 1785 were applied to each dataset. Objects with an area ≤50μm^2^ were omitted by filtering and sphericity measurements were performed on remaining surfaces.

### Statistical analysis

Biological replicates for all RNA-seq and qRT-PCR datasets represent the number of times each experiment was performed independently using pooled larvae. Whereas biological replicates for all imaging datasets represent the number of individual larvae analyzed for each experiment. Technical replicates represent repeated measurements from the same biological replicate/sample and were only used for qRT-PCR analyses to account for well-to-well variation. All statistical analyses and graphical representation of raw data were performed using Prism 8 (GraphPad Software, San Diego, California). Normal (Gaussian) distribution was assessed using the D’Agostino-Pearson omnibus normality test. In datasets with a normal distribution, significance was determined using the unpaired t-test with Welch’s correction; in datasets with non-normal distribution, nonparametric Mann-Whitney tests were performed to calculate p-values. For all graphical data presentation, the median represents the measure of center. Exact p-values, n values, number of experiments, and statistical analyses for each data comparison can be found in Table 2.

**Table 2.**
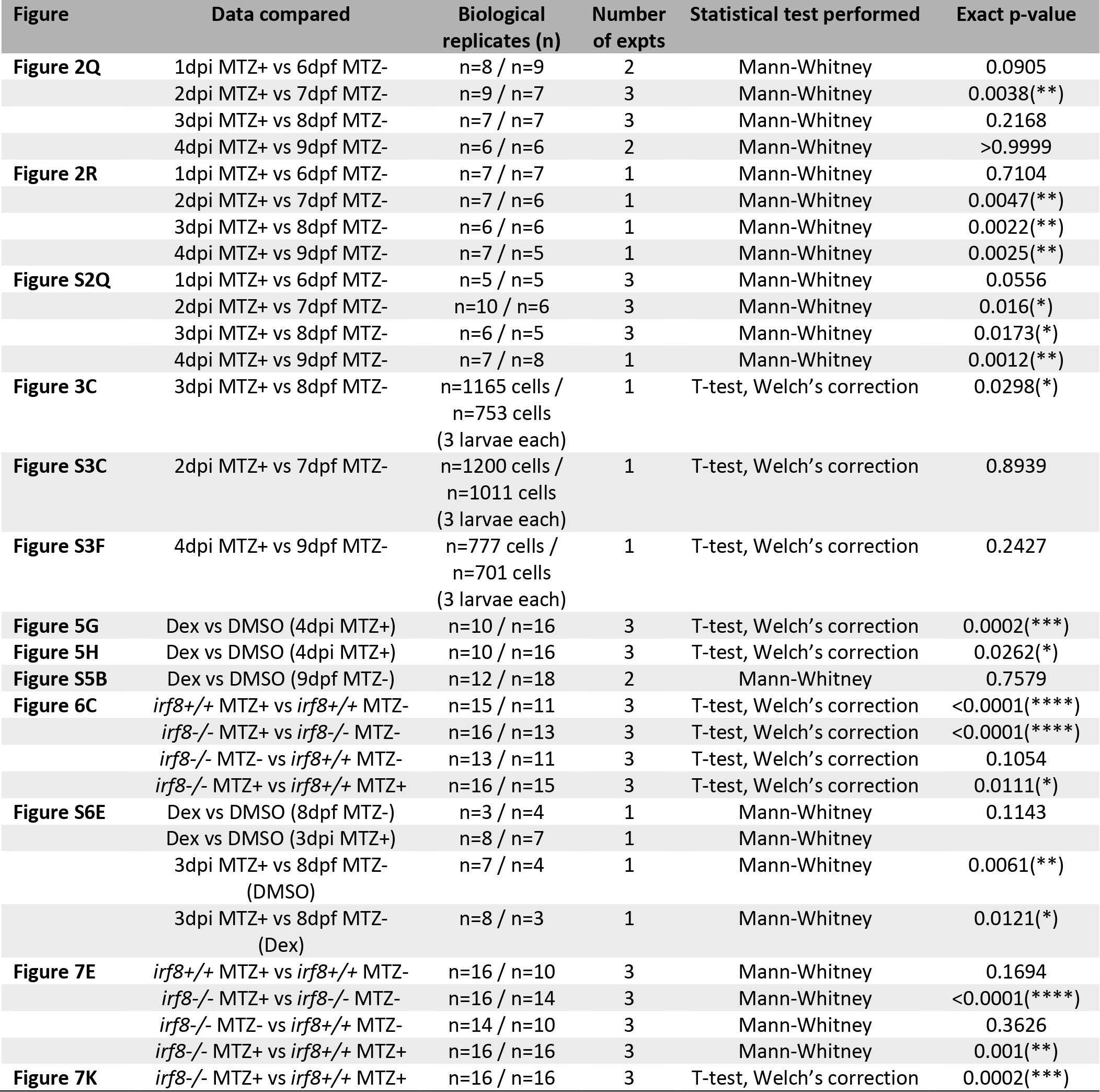
Statistical information organized by figure.

## ACKNOWLEDGEMENTS

The work described herein was supported by the National Institutes of Health (T32-EY17271 to L.L.L., RO1-EY29410 to J.M.G, and NIH CORE Grant P30-EY08098 to the Department of Ophthalmology); the UPMC Immune Transplant & Therapy Center (to L.L.L. and J.M.G.); the Pennsylvania Lions Sight Conservation and Eye Research Foundation (to L.L.L. and J.M.G.); the Whitman Center at the Marine Biological Laboratory (to J.M.G.); the Charles and Louella Snyder Retinal Regeneration Fund (to J.M.G.); the Macular Degeneration Research Program of the BrightFocus Foundation (M2016067 to J.M.G); and the E. Ronald Salvitti Chair in Ophthalmology Research (to J.M.G.). Additional support was received from the Martha Wandrisco Neff Research Award in Macular Degeneration (to L.L.L. and N.J.H.), an Honors College HEAL Research Fellowship (to S.M.G), the Eye & Ear Foundation of Pittsburgh, and an unrestricted grant from Research to Prevent Blindness, New York, NY. We also wish to thank Dr. G. Burch Fisher III for statistical discussion and expertise, Veronica Spector for editorial assistance, Dr. Hugh Hammer for expert zebrafish care, Drs. Paul Maddox and Joel Smith of Mizar Imaging for advice on light-sheet microscopy, and Dr. Marko Horb and members of his laboratory for hosting J.M.G and L.L.L. while at the Marine Biological Laboratory for light-sheet imaging experiments.

