## Supplemental Material for "The immune response is a critical regulator of zebrafish retinal pigment epithelium regeneration"

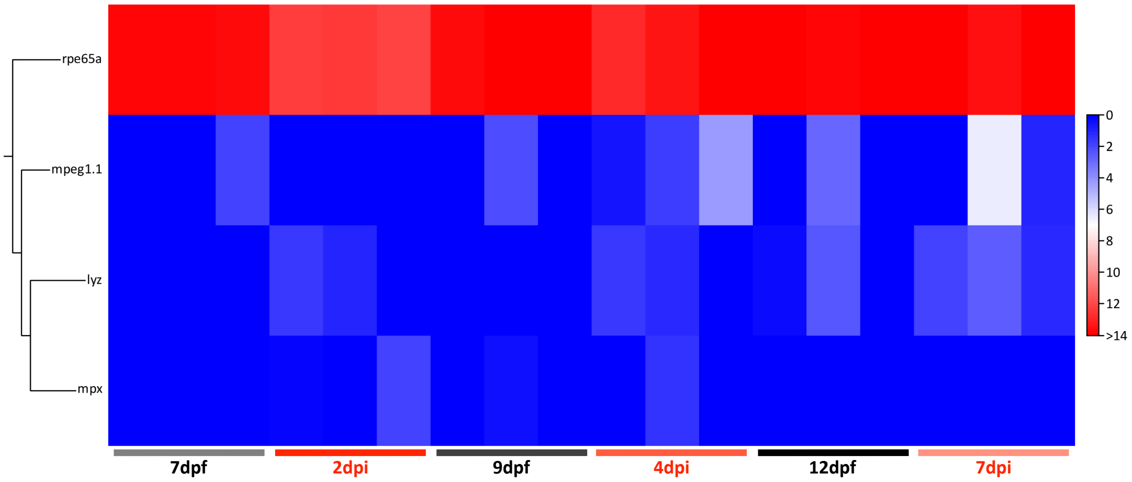


**Figure S1. *rpe65a* is highly expressed in FACS-isolated eGFP$^+$ cell populations.** Heatmap showing hierarchical clustering of RPE *(rpe65a)* and leukocyte *(mpeg1.1, mpx, lyz)* marker genes in eGFP$^+$ FACS-purified cells from *rpe65a:*nfsB-eGFP MTZ- and MTZ+ treated larvae. *rpe65a* is highly expressed whereas leukocyte markers are low or not expressed across all timepoints. Heatmap legend represents $\log_2$(TPM+1). Definitions as follows: dpf, days post-fertilization; dpi, days post-injury; FACS, fluorescence-activated cell sorting; MTZ, metronidazole; RPE, retinal pigment epithelium; TPM, transcripts per million.

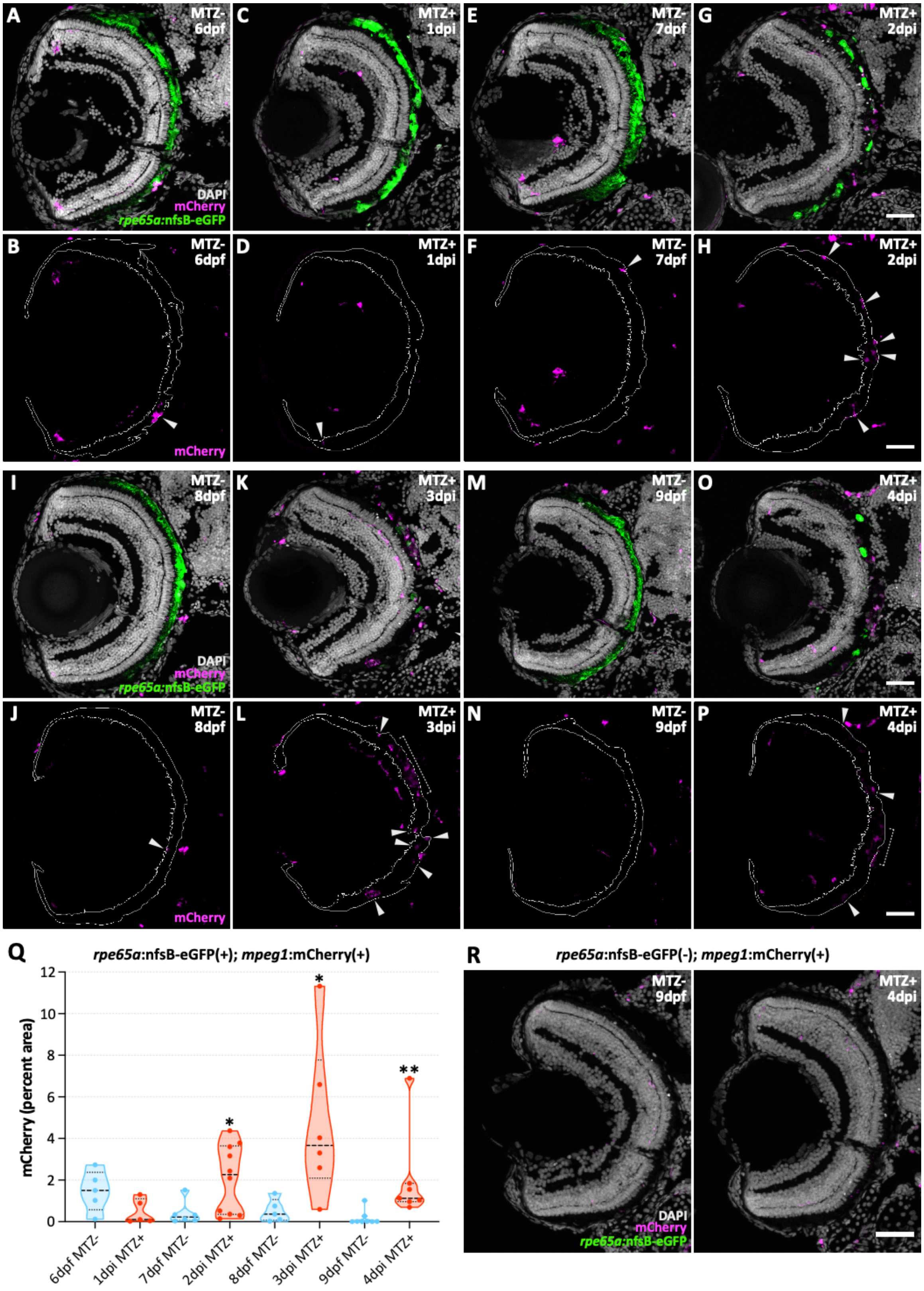


**Figure S2. Macrophages/microglia infiltrate the RPE injury site during regeneration. (A-P)** Fluorescent confocal micrographs of transverse sections from MTZ- unablated and MTZ+ ablated Tg(*mpeg1:*mCherry;

*rpe65a:*nfsB-eGFP) larval eyes from 6dpf/1dpi to 9dpf/4dpi. Images show progressive influx of MΦs/µglia labeled with mCherry antibody *(magenta)*, beginning at 2dpi **(G,H)** and peaking at 3dpi **(K,L)**. In mCherry-only panels, *white arrowheads* point to mCherry$^+$ cells within the RPE layer *(white lines)* and *white dashed brackets* **(L,P)** indicate mCherry$^+$ clusters where single cells are difficult to distinguish. **(Q)** Violin plots showing quantification of the percent area occupied by mCherry$^+$ staining during RPE regeneration. Data show significant increases in MTZ+ ablated larvae at 2dpi (p=0.016), 3dpi (p=0.0173), and 4dpi (p=0.0012) when compared to age-matched MTZ- unablated controls. *Dashed black lines* represent the median and *dotted black lines* represent quartiles. Statistics (number of experiments, biological replicates (n), statistical test, and p-values) can be found in Table 2. **(R)** Representative fluorescent confocal micrographs of transverse sections from 9dpf untreated (MTZ-; n=6 from 2 experiments) and 4dpi MTZ-treated (n=8 from 2 experiments) Tg(*mpeg1:*mCherry) larval eyes. Images show mCherry$^+$ cells do not infiltrate the RPE layer in larvae lacking the *rpe65a:*nfsB-eGFP transgene. For all micrographs, *white* (DAPI) labels nuclei, *magenta* labels mCherry$^+$ cells, and *green* labels endogenous *rpe65a:*nfsB-eGFP. Scale bars represent 40µm. Dorsal is up; definitions as follows: *, p-value ≤0.05; **, p-value ≤0.01; dpf, days post-fertilization; dpi, days post-injury; MΦs/µglia, macrophages/microglia; MTZ, metronidazole; RPE, retinal pigment epithelium.

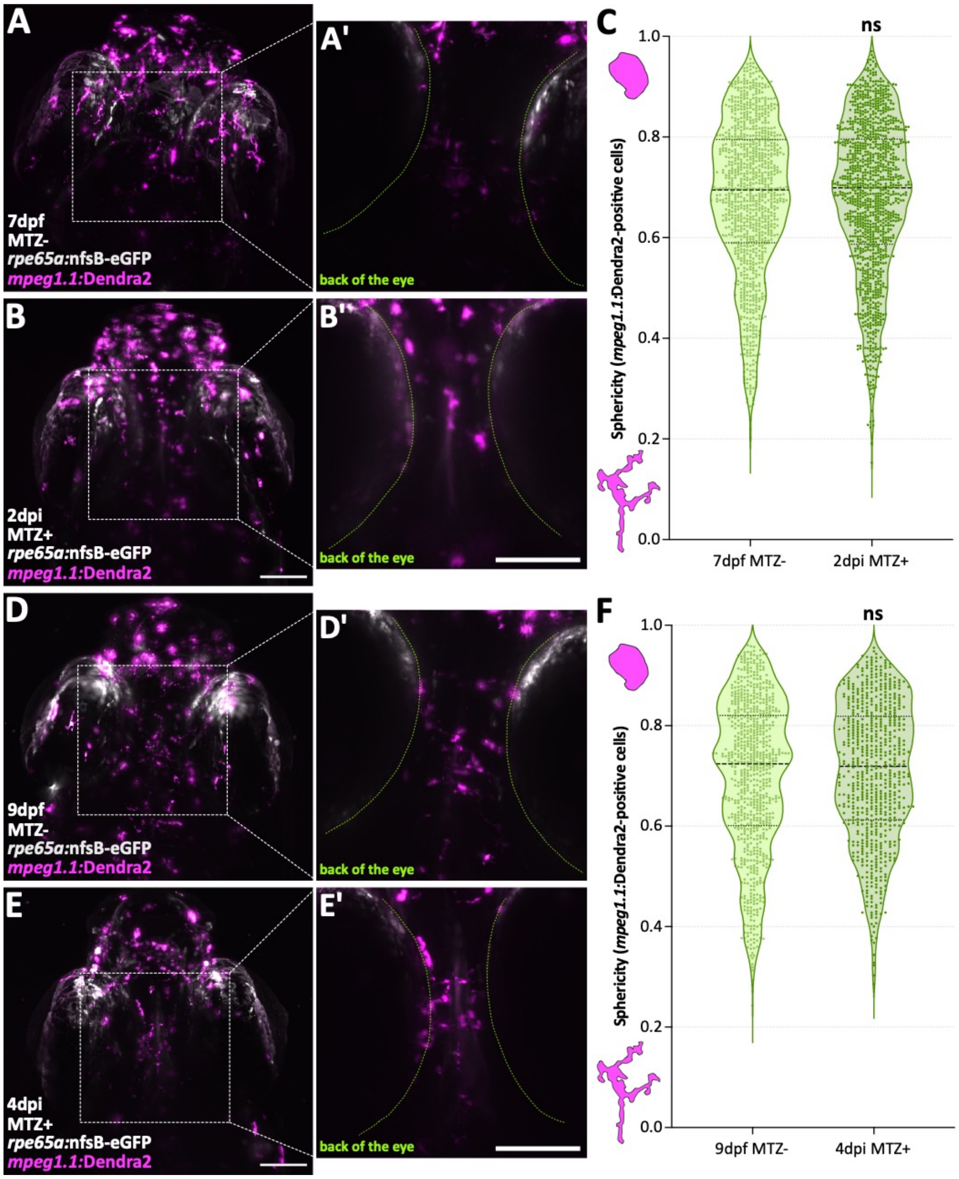


**Figure S3. Anterior macrophages/microglia show no differences in sphericity at 2 and 4 days post-RPE ablation.** *In vivo* fluorescent light-sheet micrographs from 7dpf MTZ- unablated **(A,A')** and 2dpi MTZ+ ablated **(B,B')** Tg(*mpeg1.1:*Dendra2; *rpe65a:*nfsB-eGFP) whole larvae. **(A',B')** Digital zooms of cropped 100µm z-stacks (z-step 100-200) showing the back of the eye *(green dashed line)*. **(C)** Violin plots showing no significant difference in the sphericity of *mpeg1.1:*Dendra2(red)$^{+}$ cells in the anterior portion of zebrafish larvae at 2dpi compared to 7dpf (p=0.8939). Data represent n=1011 cells from 3 larvae (MTZ-) and n=1200 cells from 3 larvae (MTZ+). **(D-E')** *In vivo* fluorescent light-sheet micrographs from 9dpf MTZ- **(D,D')** and 4dpi MTZ+ **(E,E')** Tg(*mpeg1.1:*Dendra2; *rpe65a:*nfsB-eGFP) whole larvae. **(D',E')** Digital zooms of cropped 100µm z-stacks (z-step 100-200) showing the back of the eye *(green dashed line)*. **(F)** Violin plots showing no significant difference in cell sphericity at 4dpi compared to 9dpf (p=0.2427). Data represent n=777 cells from 3 larvae (MTZ-) and n=701 cells from 3 larvae (MTZ+). **(A-B'; D-E')** *White* labels endogenous *rpe65a:*nfsB-eGFP and *magenta* labels endogenous *mpeg1.1:*Dendra2 (red). All scale bars represent 100µm. **(C,F)** *Dashed black lines* represent the median and *dotted black lines* represent quartiles. Statistics (number of experiments, biological replicates (n), statistical test, and p-values) can be found in Table 2. Anterior is up; definitions as follows: dpf, days post-fertilization; dpi, days post-injury; MTZ, metronidazole; ns, not significant; RPE, retinal pigment epithelium.

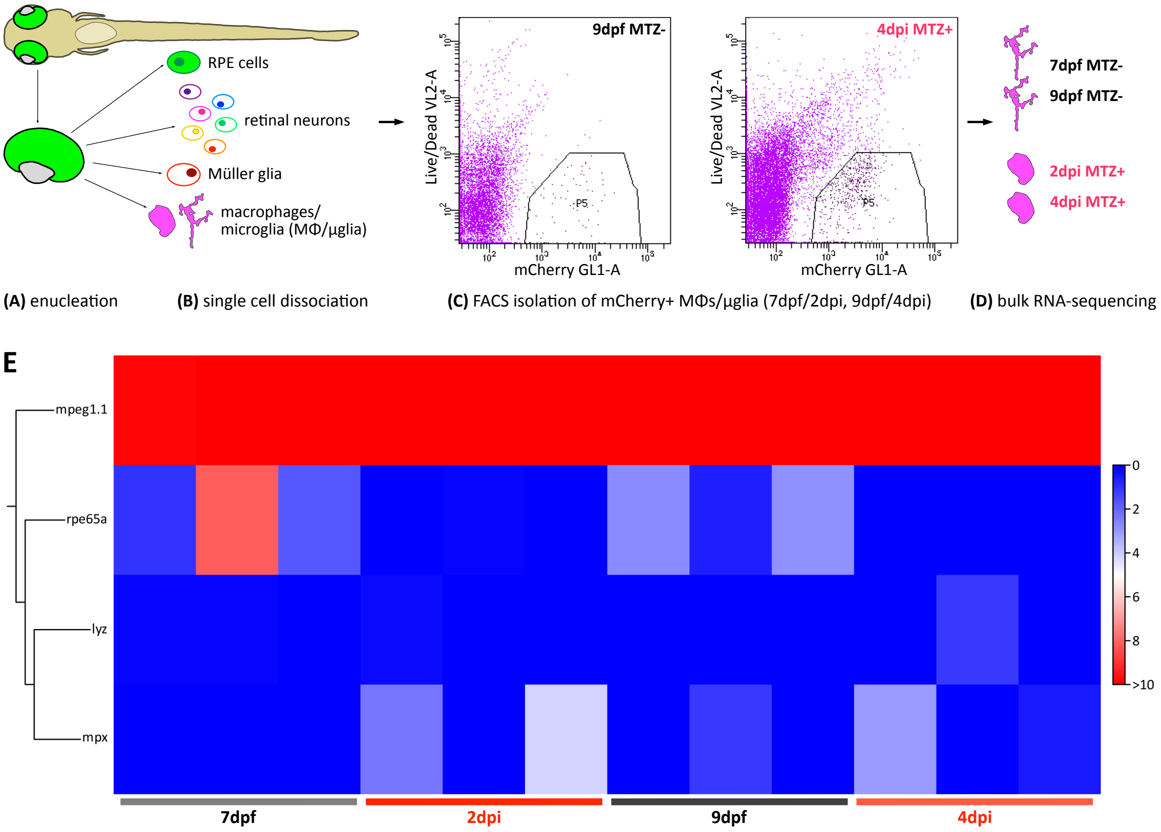


**Figure S4. Experimental workflow for isolating mCherry$^+$ macrophages/microglia from larval zebrafish.** Schematic showing enucleation **(A)**, dissociation **(B)**, and isolation of mCherry$^+$ MΦs/µglia by FACS for bulk RNA-sequencing at two timepoints post-RPE ablation **(C,D)**. Example 9dpf and 4dpi FACS plots show gate set for cell sorting (**C**; *P5*). There are more mCherry$^+$ events present in the P5 gate in ablated larvae. **(E)** Heatmap showing hierarchical clustering of leukocyte *(mpeg1.1, mpx, lyz)* and RPE *(rpe65a)* marker genes in mCherry$^+$ FACS-purified cells from Tg(*rpe65a:*nfsB-eGFP;*mpeg1*:mCherry) MTZ- and MTZ+ treated larvae. *mpeg1.1* is highly expressed whereas neutrophil *(mpx, lyz)* and RPE *(rpe65a)* markers are low or not expressed across all timepoints with the exception of one replicate at 7dpf. Heatmap legend represents $\log_2$(TPM+1). Definitions as follows: dpf, days post-fertilization; dpi, days post-injury; FACS, fluorescence-activated cell sorting; MΦs/µglia, macrophages/microglia; MTZ, metronidazole; RPE, retinal pigment epithelium; TPM, transcripts per million.

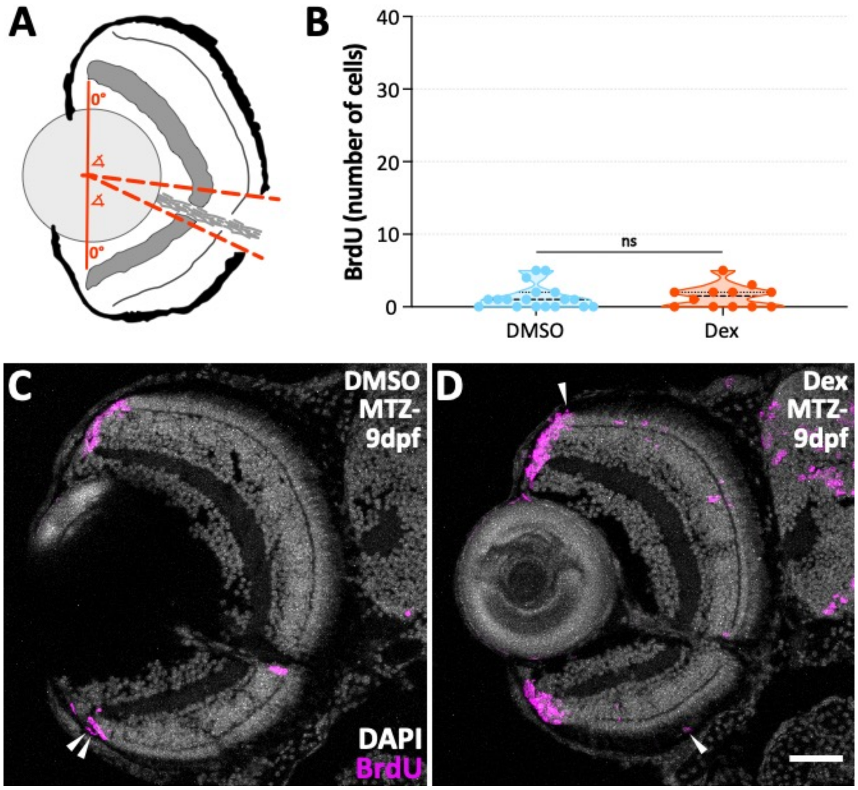


**Figure S5. Dexamethasone does not affect RPE proliferation in unablated larvae. (A)** Schematic showing quantification of RPE regeneration based on pigment recovery. A dorsal-ventral line with midpoint *(solid red line)* was drawn and dorsal and ventral angle measurements were made between 0° and the point where continuous pigmentation stopped *(dashed red lines)*. **(B)** Violin plots showing no significant difference in the number of BrdU$^+$ cells in dexamethasone-treated larvae when compared to DMSO controls (p=0.7579). *Dashed black lines* represent the median and *dotted black lines* represent quartiles. Statistics (number of experiments, biological replicates (n), statistical test, and p-values) can be found in Table 2. **(C,D)** Fluorescent confocal micrographs of transverse sections from 9dpf MTZ- unablated DMSO- **(C)** and dexamethasone-treated **(D)** Tg(*rpe65a:*nfsB-eGFP) larval eyes. Images show few BrdU$^+$ cells in these larvae. *White arrowheads* highlight BrdU-labeled cells *(magenta)* and *white* (DAPI) labels nuclei. Scale bar represents 40µm. Dorsal is up; definitions as follows: BrdU, bromodeoxyuridine; dex, dexamethasone; DMSO, dimethyl sulfoxide; MTZ, metronidazole; ns, not significant; RPE, retinal pigment epithelium.

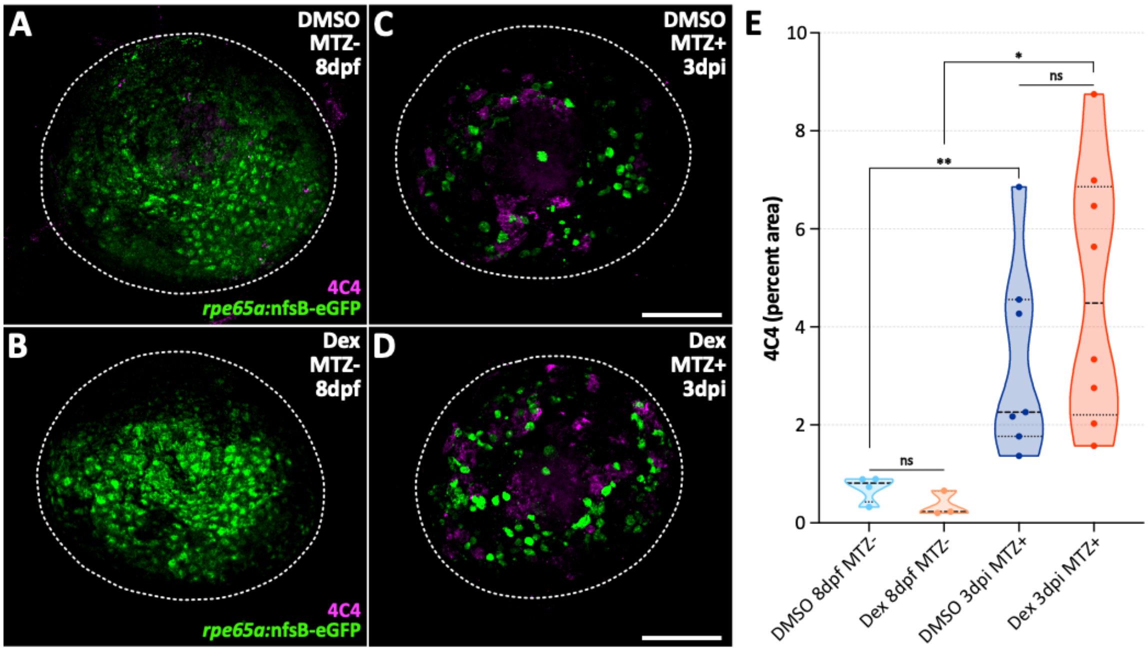


**Figure S6. Dexamethasone does not affect macrophage/microglia infiltration post-RPE ablation.** Fluorescent confocal micrographs of 8dpf MTZ- unablated **(A,B)** and 3dpi MTZ+ ablated **(C,D)** Tg(*rpe65a:*nfsB-eGFP) whole mount eyes labeled with 4C4 antibody to mark MΦs/μglia. Larvae were treated either with 0.05% DMSO (**A,C**; vehicle control) or 50μM dexamethasone **(B,D)** from 4-8dpf. Images show clear ablation of the RPE and infiltration of 4C4⁺ cells post-injury in both DMSO- and dexamethasone-treated larvae. *White dashed lines* designate edge of whole mount larval eye. *Magenta* labels 4C4 and *green* labels endogenous *rpe65a:*nfsB-eGFP. Scale bars represent 100μm. **(E)** Violin plots showing quantification of the percent area occupied by 4C4⁺ staining during RPE regeneration. Data show significant increases by treatment in MTZ+ ablated larvae when compared to age-matched MTZ- unablated controls (DMSO, p=0.0061; Dex, p=0.0121), but no significant different between treatments in MTZ- unablated (p=0.1143) or MTZ+ ablated (p=0.3357) larvae. *Dashed black lines* represent the median and *dotted black lines* represent quartiles. Statistics (number of experiments, biological replicates (n), statistical test, and p-values) can be found in Table 2. Dorsal is up; definitions as follows: *, p-value ≤0.05; **, p-value ≤0.01; dex, dexamethasone; DMSO, dimethyl sulfoxide; dpf, days post-fertilization; dpi, days post-injury; ns, not significant; MΦs/μglia, macrophages/microglia; MTZ, metronidazole; RPE, retinal pigment epithelium.

**Table S1.** Top 50 upregulated genes from RPE (eGFP+) cells at 2dpi compared to 7dpf.

| Gene name (1-25) | Fold change | P-value | FDR p-value | Gene name (26-50) | Fold change | P-value | FDR p-value |
|---|---|---|---|---|---|---|---|
| *il11b* | 35,950.23 | 1.83E-10 | 8.25E-08 | *CU571324.1* | 39.96 | 1.67E-09 | 5.69E-07 |
| *edn2* | 5,906.55 | 1.27E-07 | 2.36E-05 | *rgs5b* | 39.31 | 3.95E-12 | 2.71E-09 |
| *nppb* | 2,386.58 | 7.14E-11 | 3.66E-08 | *BX088646.1* | 36.15 | 1.74E-12 | 1.3E-09 |
| *saa* | 2,303.71 | 5.78E-06 | 6.05E-04 | *cbx7a* | 30.57 | 6.49E-12 | 4.12E-09 |
| *dkk1a* | 1,579.88 | 1.55E-14 | 2.02E-11 | *helq* | 30.43 | 6.37E-14 | 7.29E-11 |
| *lepb* | 1,312.82 | 2.37E-05 | 1.83E-03 | *lye* | 29.34 | 5.28E-09 | 1.62E-06 |
| *il34* | 1,200.32 | 3.83E-13 | 3.37E-10 | *ptgdsb.1* | 29.33 | 7.65E-10 | 2.95E-07 |
| *CR762483.1* | 917.39 | 1.35E-10 | 6.41E-08 | *npy* | 27.08 | 4.84E-08 | 1.03E-05 |
| *plekhs1* | 257.08 | 2.8E-12 | 2E-09 | *AL954191.1* | 25.76 | 1.07E-06 | 1.44E-04 |
| *CABZ01012857.1* | 232.24 | 1.44E-15 | 2.61E-12 | *mid1ip1a* | 25.14 | 7.8E-08 | 1.53E-05 |
| *ccl34a.4* | 209.43 | 2.2E-06 | 2.62E-04 | *si:dkey-40h20.1* | 24.11 | 1.54E-08 | 4.29E-06 |
| *scpp8* | 147.01 | 1.25E-11 | 7.65E-09 | *dusp8b* | 23.77 | 5.57E-13 | 4.55E-10 |
| *cxcl8a* | 123.89 | 2.37E-11 | 1.31E-08 | *BX470240.1* | 23.51 | 1.62E-08 | 4.44E-06 |
| *si:busm1-57f23.1* | 89.57 | 1.11E-16 | 2.38E-13 | *nfkbiaa* | 23.05 | 5.44E-12 | 3.52E-09 |
| *CR925709.2* | 66.72 | 1.03E-09 | 3.88E-07 | *il4r.1* | 22.59 | 2.11E-11 | 1.21E-08 |
| *mmp13b* | 58.02 | 9.77E-15 | 1.34E-11 | *nmu* | 22.28 | 2.43E-06 | 2.81E-04 |
| *cidec* | 51.78 | 6.66E-15 | 9.94E-12 | *tubb6* | 22.17 | 6.32E-10 | 2.49E-07 |
| *tmem238b* | 46.99 | 7.39E-08 | 1.46E-05 | *igfbp1a* | 21.28 | 7.96E-11 | 3.96E-08 |
| *zgc:198419* | 45.46 | 2.38E-11 | 1.31E-08 | *smarcd2* | 21.22 | 8.62E-08 | 1.66E-05 |
| *mns1* | 45.40 | 1.59E-14 | 2.02E-11 | *hsd17b12a* | 20.98 | 1.48E-09 | 5.31E-07 |
| *cdh1* | 45.28 | 3.03E-11 | 1.6E-08 | *casq2* | 19.82 | 9.35E-09 | 2.7E-06 |
| *cfl1l* | 43.07 | 1.16E-10 | 5.59E-08 | *si:dkey-184p18.2* | 18.96 | 7.14E-07 | 1.06E-04 |
| *BX005392.3* | 42.29 | 2.57E-11 | 1.4E-08 | *mcamb* | 18.95 | 3.23E-07 | 5.49E-05 |
| *cxcl18b* | 42.23 | 3.21E-13 | 2.9E-10 | *rbms1a* | 18.74 | 8.58E-12 | 5.35E-09 |
| *isg15* | 40.84 | 3.05E-13 | 2.83E-10 | *AL954861.1* | 18.21 | 4.43E-10 | 1.84E-07 |

Filtering parameters: mean RPKM ≥5, fold change ≥2, false discovery rate (FDR) p-value ≤0.05.

**Table S2.** Top 50 upregulated genes from RPE (eGFP$^+$) cells at 4dpi compared to 9dpf.

| Gene name (1-25) | Fold change | P-value | FDR p-value | Gene name (26-50) | Fold change | P-value | FDR p-value |
|---|---|---|---|---|---|---|---|
| *CU927934.1* | 2,385.11 | 2.22E-16 | 2.72E-13 | *epcam* | 16.42 | 3.82E-04 | 0.02 |
| *nppb* | 1,008.65 | 3.91E-09 | 1.11E-06 | *clic2* | 16.16 | 1.31E-06 | 1.91E-04 |
| *dkk1a* | 670.56 | 1.89E-15 | 2.02E-12 | *cxcl8a* | 15.91 | 9.24E-06 | 9.62E-04 |
| *lye* | 131.45 | 1.86E-13 | 1.42E-10 | *hpse* | 15.80 | 2.43E-13 | 1.81E-10 |
| *cyp2k22* | 92.24 | 1.7E-12 | 9.72E-10 | *phlda2* | 15.79 | 7.99E-10 | 2.69E-07 |
| *si:ch73-6k14.2* | 82.83 | 1.11E-16 | 1.41E-13 | *apoeb* | 15.31 | 8.4E-09 | 2.24E-06 |
| *ccl34a.4* | 77.08 | 4.62E-08 | 1.04E-05 | *timp4.1* | 15.30 | 1.91E-09 | 5.9E-07 |
| *si:dkey-23a13.2* | 64.19 | 4.44E-16 | 5.26E-13 | *grn1* | 15.26 | 4.61E-05 | 3.56E-03 |
| *si:busm1-57f23.1* | 58.91 | 6.66E-16 | 7.62E-13 | *pdgfaa* | 14.94 | 2.22E-15 | 2.31E-12 |
| *slc43a3b* | 54.06 | 2.24E-07 | 4.05E-05 | *tes* | 14.84 | 2.51E-07 | 4.47E-05 |
| *mmp9* | 52.49 | 1.91E-10 | 7.91E-08 | *tagln2* | 14.46 | 2.83E-06 | 3.62E-04 |
| *cart3* | 50.74 | 2.53E-06 | 3.29E-04 | *ptgdsb.1* | 14.25 | 1.53E-05 | 1.47E-03 |
| *fosl1b* | 48.20 | 2.26E-10 | 9.24E-08 | *anxa2a* | 14.09 | 8.44E-12 | 4.53E-09 |
| *ptgs2a* | 46.48 | 4.29E-08 | 9.82E-06 | *irf8* | 14.01 | 1.38E-07 | 2.64E-05 |
| *nmu* | 43.52 | 3.11E-09 | 9.05E-07 | *nat8l* | 13.99 | 1.02E-09 | 3.39E-07 |
| *ctss2.1* | 36.37 | 4.6E-06 | 5.41E-04 | *si:ch73-160h15.3* | 13.98 | 3.72E-04 | 0.02 |
| *zgc:158343* | 30.36 | 7.76E-09 | 2.08E-06 | *arpp21* | 13.87 | 1.14E-07 | 2.24E-05 |
| *edn2* | 30.26 | 2.63E-07 | 4.64E-05 | *cxcl14* | 13.56 | 6.78E-10 | 2.33E-07 |
| *cd99* | 27.03 | 1.7E-12 | 9.72E-10 | *atp1a3b* | 12.99 | 1.7E-12 | 9.72E-10 |
| *acp6* | 26.81 | 3.39E-14 | 3.14E-11 | *scpp8* | 12.81 | 1.83E-06 | 2.54E-04 |
| *il34* | 25.10 | 5.72E-10 | 2E-07 | *si:ch211-105c13.3* | 12.57 | 1.39E-06 | 2.01E-04 |
| *mcamb* | 19.44 | 1.8E-08 | 4.4E-06 | *rac2* | 12.49 | 2.41E-05 | 2.11E-03 |
| *stm* | 17.37 | 1.46E-07 | 2.75E-05 | *cpa2* | 12.09 | 1.6E-05 | 1.51E-03 |
| *CU984579.1* | 17.16 | 8.33E-15 | 8.17E-12 | *creld2* | 11.87 | 1.71E-08 | 4.26E-06 |
| *ctsk* | 16.71 | 4.14E-10 | 1.53E-07 | *mt2* | 11.55 | 5.75E-08 | 1.23E-05 |

Filtering parameters: mean RPKM ≥5, fold change ≥2, false discovery rate (FDR) p-value ≤0.05.

**Table S3.** Top upregulated genes from RPE (eGFP⁺) cells at 7dpi compared to 12dpf.

| Gene name (only 15 total after filtering) | Fold change | P-value | FDR p-value |
|---|---|---|---|
| *fshb* | 5,099.35 | 4.35E-06 | 0.01 |
| *havcr2* | 3,120.74 | 1.97E-05 | 0.03 |
| *BX649485.4* | 2,270.58 | 4.06E-05 | 0.04 |
| *stm* | 112.06 | 7.77E-16 | 2.67E-11 |
| *lye* | 98.18 | 4.12E-06 | 0.01 |
| *cfl1l* | 43.38 | 1.06E-08 | 7.3E-05 |
| *kpna2* | 38.38 | 3.04E-07 | 1.74E-03 |
| *podxl* | 33.18 | 3.41E-05 | 0.04 |
| *isg15* | 21.36 | 6.08E-06 | 0.01 |
| *thy1* | 11.36 | 1.12E-06 | 3.83E-03 |
| *ctsk* | 10.83 | 5.4E-06 | 0.01 |
| *marcksl1a* | 9.58 | 5.98E-07 | 2.57E-03 |
| *zgc:198419* | 8.88 | 7.37E-06 | 0.01 |
| *s100a10b* | 7.76 | 7.03E-06 | 0.01 |
| *si:ch73-158p21.3* | 7.75 | 5.42E-05 | 0.05 |

Filtering parameters: mean RPKM ≥5, fold change ≥2, false discovery rate (FDR) p-value ≤0.05.

**Table S4.** Top 50 downregulated genes from RPE (eGFP$^+$) cells at 2dpi compared to 7dpf.

| Gene name (1-25) | Fold change | P-value | FDR p-value | Gene name (26-50) | Fold change | P-value | FDR p-value |
|---|---|---|---|---|---|---|---|
| *pvalb4* | -8347.95 | 3.28E-06 | 0.000369 | *hes2.2* | -73.6779 | 4.05E-12 | 2.72E-09 |
| *tnni2a.4* | -5117.54 | 9.59E-06 | 0.000892 | *ch25hl1.1* | -72.3766 | 1.58E-09 | 5.51E-07 |
| *pvalb1* | -2797.35 | 2.33E-05 | 0.001804 | *krt4* | -58.5131 | 1.03E-05 | 0.000936 |
| *tpma* | -2119.11 | 2.64E-13 | 2.52E-10 | *zgc:153405* | -46.7638 | 5.49E-06 | 0.000582 |
| *mylpfa* | -2068.25 | 5.57E-11 | 2.9E-08 | *ascl1a* | -44.4477 | 1.05E-13 | 1.12E-10 |
| *ckma* | -1754.27 | 1.88E-09 | 6.19E-07 | *CU467822.1* | -42.4573 | 8.65E-10 | 3.3E-07 |
| *si:ch211-193l2.7* | -769.289 | 6.44E-15 | 9.94E-12 | *CR848784.5* | -41.1808 | 2.95E-08 | 7.09E-06 |
| *actc1b* | -600.373 | 1.74E-10 | 7.94E-08 | *opn1sw1* | -40.4664 | 3.87E-08 | 8.64E-06 |
| *cxcr4b* | -461.435 | 2.29E-14 | 2.76E-11 | *crabp2a* | -36.6931 | 8.55E-15 | 1.22E-11 |
| *her12* | -457.29 | 2.33E-14 | 2.76E-11 | *BX465834.1* | -34.5499 | 3.4E-09 | 1.08E-06 |
| *si:ch73-21g5.7* | -400.366 | 4.53E-08 | 9.67E-06 | *zgc:112234_9* | -34.1586 | 1.17E-13 | 1.22E-10 |
| *pvalb2* | -375.199 | 1.79E-06 | 0.00022 | *dao.3* | -33.9566 | 7.41E-14 | 8.2E-11 |
| *ckmb* | -274.546 | 4.81E-09 | 1.49E-06 | *tcap* | -32.8645 | 0.000205 | 0.009938 |
| *si:ch211-193l2.4* | -236.093 | 3.55E-15 | 5.99E-12 | *gadd45gb.1* | -32.1544 | 1.59E-09 | 5.51E-07 |
| *AL929185.2* | -215.612 | 2.31E-13 | 2.26E-10 | *si:ch211-113a14.18* | -31.3796 | 1.56E-06 | 0.000196 |
| *tnnc2* | -202.228 | 3.58E-08 | 8.14E-06 | *tfpia* | -31.1528 | 5.42E-13 | 4.54E-10 |
| *myl1* | -149.022 | 9.03E-09 | 2.63E-06 | *si:ch211-113a14.24* | -30.165 | 7.3E-05 | 0.004466 |
| *AL929185.1* | -147.128 | 4.45E-13 | 3.82E-10 | *aplnra* | -28.256 | 7.98E-08 | 1.56E-05 |
| *TMEM27* | -135.1 | 6.66E-16 | 1.35E-12 | *mylz3* | -28.0453 | 1.69E-05 | 0.001378 |
| *opn1mw1* | -98.4783 | 7.91E-11 | 3.96E-08 | *neurod4* | -27.5696 | 2.6E-10 | 1.13E-07 |
| *si:ch211-193l2.5* | -91.1578 | 1.22E-09 | 4.45E-07 | *zgc:173552_5* | -26.3851 | 0.000251 | 0.011513 |
| *si:ch73-330k17.3* | -85.5907 | 1.27E-11 | 7.65E-09 | *si:ch73-389b16.1* | -25.2493 | 0.000162 | 0.008375 |
| *her15.1_1* | -84.0241 | 3.66E-15 | 5.99E-12 | *si:dkey-238o13.4* | -24.5866 | 1.57E-10 | 7.3E-08 |
| *krtt1c19e* | -79.1791 | 1.76E-05 | 0.001429 | *aldob* | -23.9921 | 2.61E-12 | 1.9E-09 |
| *nme2b.2* | -78.765 | 2.32E-08 | 5.98E-06 | *lmo1* | -23.2547 | 5.72E-10 | 2.3E-07 |

Filtering parameters: mean RPKM ≥5, fold change ≤2, false discovery rate (FDR) p-value ≤0.05.

**Table S5.** Top 50 downregulated genes from RPE (eGFP⁺) cells at 4dpi compared to 9dpf.

| Gene name (1-25) | Fold change | P-value | FDR p-value | Gene name (26-50) | Fold change | P-value | FDR p-value |
|---|---|---|---|---|---|---|---|
| *CU464134.1* | -200.276 | 1.54E-10 | 6.77E-08 | *znfl2a* | -10.1748 | 7.34E-05 | 0.005143 |
| *crygmx* | -131.323 | 1.12E-06 | 0.000166 | *zic3* | -10.1067 | 1.46E-09 | 4.68E-07 |
| *hes2.2* | -72.76 | 1.67E-10 | 6.98E-08 | *ypel3* | -10.0312 | 1.19E-12 | 7.45E-10 |
| *crybb1* | -64.3886 | 4.7E-09 | 1.29E-06 | *crhbp* | -9.78379 | 5.19E-07 | 8.48E-05 |
| *cav2* | -47.233 | 1.26E-10 | 5.61E-08 | *chadlb* | -9.76533 | 2.14E-06 | 0.000287 |
| *cryba1l1* | -43.2732 | 4.67E-05 | 0.003594 | *krt97* | -8.82446 | 4.76E-11 | 2.37E-08 |
| *cryba4* | -37.6295 | 3.03E-06 | 0.000382 | *mmel1* | -8.81027 | 1.64E-09 | 5.11E-07 |
| *myoz1b* | -25.5932 | 3.69E-10 | 1.39E-07 | *svila* | -8.67118 | 1.42E-13 | 1.13E-10 |
| *crygn2* | -23.9837 | 7.62E-05 | 0.005273 | *TMEM269* | -8.53631 | 5.16E-07 | 8.48E-05 |
| *zgc:171686* | -23.9712 | 2.58E-11 | 1.32E-08 | *phyhipla* | -8.22502 | 5.61E-13 | 3.77E-10 |
| *si:ch211-270g19.5* | -23.5055 | 1.05E-09 | 3.48E-07 | *aldh1l1* | -7.92923 | 1.31E-09 | 4.24E-07 |
| *slc22a6l* | -23.0486 | 1.07E-12 | 6.8E-10 | *zgc:114046* | -7.80873 | 4.39E-06 | 0.000519 |
| *CR848784.5* | -20.9448 | 2.5E-09 | 7.53E-07 | *jam2a* | -7.7691 | 4.19E-13 | 2.93E-10 |
| *igf1* | -20.4321 | 1.42E-05 | 0.001388 | *zgc:194659* | -7.75339 | 1.31E-07 | 2.51E-05 |
| *ambp* | -18.8113 | 1.92E-12 | 1.07E-09 | *col9a1a* | -7.74794 | 3.34E-11 | 1.68E-08 |
| *her4.3* | -18.1182 | 1.44E-05 | 0.001394 | *vasna* | -7.66627 | 9.81E-12 | 5.18E-09 |
| *htra1b* | -17.0303 | 1.56E-10 | 6.8E-08 | *acbd7* | -7.63206 | 1.61E-09 | 5.06E-07 |
| *slc43a2b* | -15.9854 | 2.89E-15 | 2.91E-12 | *zgc:194209* | -7.47792 | 5.04E-08 | 1.12E-05 |
| *si:ch211-193l2.4* | -15.5122 | 0.000105 | 0.006775 | *zgc:112234_9* | -7.43733 | 0.000305 | 0.01525 |
| *si:ch211-193l2.3* | -14.3345 | 2.74E-05 | 0.002303 | *lpar6b* | -7.30038 | 4.3E-12 | 2.34E-09 |
| *zgc:154164* | -13.9562 | 8.08E-05 | 0.005547 | *pcxb* | -7.2664 | 1.61E-06 | 0.000229 |
| *zdhhc15a* | -13.6603 | 1.33E-08 | 3.42E-06 | *gng3* | -7.03947 | 0.000631 | 0.026321 |
| *tmem38a* | -12.4258 | 4.72E-06 | 0.000551 | *slc13a4* | -6.98471 | 1.48E-11 | 7.71E-09 |
| *insm1a* | -10.9031 | 0.000365 | 0.017584 | *inpp4ab* | -6.82072 | 5.1E-10 | 1.82E-07 |
| *si:dkeyp-120h9.1* | -10.8062 | 5.16E-06 | 0.000596 | *znf326* | -6.79041 | 5.79E-09 | 1.58E-06 |

Filtering parameters: mean RPKM ≥5, fold change ≤2, false discovery rate (FDR) p-value ≤0.05.

**Table S6.** Top downregulated genes from RPE ($eGFP^+$) cells at 7dpi compared to 12dpf.

| Gene name (only 3 total after filtering) | Fold change | P-value | FDR p-value |
|---|---|---|---|
| *wu:fj16a03* | -251.29 | 8.1E-11 | 6.95E-07 |
| *ccl25b* | -151.144 | 5.82E-06 | 0.012279 |
| *si:dkeyp-120h9.1* | -5.02068 | 4.32E-05 | 0.042339 |

Filtering parameters: mean RPKM ≥5, fold change ≤2, false discovery rate (FDR) p-value ≤0.05.

**Table S7.** Top 50 upregulated genes from macrophages/microglia (mCherry$^{+}$) cells at 2dpi compared to 7dpf.

| Gene name (1-25) | Fold change | P-value | FDR p-value | Gene name (26-50) | Fold change | P-value | FDR p-value |
|---|---|---|---|---|---|---|---|
| *si:dkey-33i11.4* | 1,478.80 | 7.4E-04 | 0.05 | *prr11* | 13.19 | 7.05E-09 | 9.17E-06 |
| *trpv6* | 960.65 | 3.24E-07 | 2.75E-04 | *timp2b* | 12.73 | 6.85E-09 | 9.17E-06 |
| *ifitm1* | 356.80 | 2.51E-08 | 2.52E-05 | *si:dkey-27h10.2* | 11.18 | 1.55E-04 | 0.02 |
| *si:ch73-343l4.8* | 335.74 | 1.67E-14 | 1.84E-10 | *mmp9* | 11.03 | 6.78E-04 | 0.05 |
| *il4* | 241.16 | 6.28E-05 | 0.01 | *CT030188.1* | 10.96 | 4.59E-07 | 3.5E-04 |
| *si:dkey-192k22.2* | 158.65 | 1.13E-05 | 3.78E-03 | *s100a10b* | 10.71 | 2.01E-05 | 5.37E-03 |
| *s100a11* | 110.03 | 6.75E-06 | 2.51E-03 | *kif20a* | 9.70 | 1.33E-05 | 4.31E-03 |
| *scpp8* | 84.96 | 2.86E-13 | 1.58E-09 | *anxa1a* | 9.59 | 9.11E-05 | 0.02 |
| *si:dkey-21e2.3_2* | 72.77 | 1.16E-04 | 0.02 | *ing5b* | 9.21 | 4.06E-04 | 0.04 |
| *anxa2a* | 70.72 | 5.13E-13 | 1.89E-09 | *si:ch211-126c2.4* | 9.04 | 5.42E-05 | 0.01 |
| *vcanb* | 67.14 | 3.7E-13 | 1.63E-09 | *f13a1b* | 8.67 | 1.27E-04 | 0.02 |
| *si:dkey-7i4.24* | 59.95 | 7.52E-07 | 5.22E-04 | *prc1b* | 8.43 | 7.32E-06 | 2.65E-03 |
| *mmp13a_1* | 58.17 | 9.28E-09 | 1.08E-05 | *nuf2* | 7.88 | 1.28E-04 | 0.02 |
| *zgc:110286* | 53.42 | 2.32E-04 | 0.03 | *ano5b* | 7.84 | 6.07E-04 | 0.04 |
| *ccnb2* | 28.56 | 3.69E-05 | 8.46E-03 | *zgc:86764* | 7.69 | 1.05E-04 | 0.02 |
| *cybrd1* | 27.96 | 6.81E-06 | 2.51E-03 | *dctd* | 7.69 | 4.93E-04 | 0.04 |
| *ccl34a.3* | 26.77 | 2.89E-04 | 0.03 | *ccnb1* | 7.36 | 1.63E-05 | 4.81E-03 |
| *cd276* | 20.25 | 4.37E-07 | 3.45E-04 | *smc2* | 7.31 | 2.02E-05 | 5.37E-03 |
| *mapk12b* | 19.54 | 4.3E-08 | 4.13E-05 | *cks2* | 7.06 | 5.95E-04 | 0.04 |
| *moxd1* | 19.27 | 2.22E-06 | 1.18E-03 | *cfbl* | 7.04 | 2.56E-04 | 0.03 |
| *cfh* | 18.42 | 2.38E-06 | 1.22E-03 | *tyms* | 6.84 | 1.18E-04 | 0.02 |
| *adam8a* | 16.79 | 2.66E-07 | 2.35E-04 | *si:dkey-66i24.9* | 6.68 | 2.04E-04 | 0.02 |
| *si:zfos-2330d3.7_2* | 16.35 | 1.53E-05 | 4.67E-03 | *ccl34a.4* | 6.67 | 5.39E-05 | 0.01 |
| *arhgef39* | 16.09 | 6.97E-09 | 9.17E-06 | *ccna2* | 6.65 | 4.7E-05 | 9.7E-03 |
| *mogat3b* | 15.27 | 3.98E-04 | 0.04 | *tagln2* | 6.64 | 1.22E-04 | 0.02 |

Filtering parameters: mean RPKM ≥5, fold change ≥2, false discovery rate (FDR) p-value ≤0.05.

**Table S8.** Top 50 upregulated genes from macrophages/microglia (mCherry+) cells at 4dpi compared to 9dpf.

| Gene name (1-25) | Fold change | P-value | FDR p-value | Gene name (26-50) | Fold change | P-value | FDR p-value |
|---|---|---|---|---|---|---|---|
| *ncaph* | 4,912.19 | 2.59E-06 | 2.31E-04 | *plk1* | 33.10 | 8.82E-10 | 3.35E-07 |
| *dctpp1* | 2,555.98 | 2.5E-05 | 1.46E-03 | *ccl34b.8* | 32.83 | 5.35E-13 | 6.93E-10 |
| *hbae3* | 2,513.11 | 1.18E-04 | 5E-03 | *cdca5* | 32.22 | 1.06E-05 | 7.39E-04 |
| *AL935210.1* | 1,654.52 | 3.18E-04 | 0.01 | *aurka* | 31.01 | 1.76E-06 | 1.72E-04 |
| *hbbe1.1* | 1,437.31 | 4.39E-04 | 0.01 | *BX323596.2* | 30.61 | 2.22E-16 | 5.43E-13 |
| *hbbe1.2* | 1,227.76 | 5.74E-04 | 0.02 | *cdc20* | 28.46 | 5.36E-08 | 1.04E-05 |
| *hbae1.1_2* | 944.12 | 7.81E-04 | 0.02 | *slc29a2* | 26.40 | 1.01E-07 | 1.75E-05 |
| *haus4* | 899.53 | 4.58E-10 | 2.06E-07 | *smc2* | 26.39 | 2.55E-10 | 1.34E-07 |
| *ccl39.1* | 300.37 | 3.52E-06 | 3.05E-04 | *si:ch73-28h20.1* | 26.13 | 2.57E-04 | 9.25E-03 |
| *cxcl8b.1* | 196.16 | 3.46E-08 | 7.47E-06 | *ndc80* | 25.57 | 6.29E-08 | 1.19E-05 |
| *dlgap5* | 85.62 | 3.15E-12 | 2.77E-09 | *zgc:194627* | 24.62 | 5.61E-07 | 6.79E-05 |
| *zgc:173552_3* | 78.63 | 8.15E-07 | 9.15E-05 | *ncapg* | 24.30 | 1.18E-06 | 1.22E-04 |
| *spc24* | 70.59 | 1.39E-09 | 4.92E-07 | *hp_2* | 24.17 | 7.45E-06 | 5.45E-04 |
| *mad2l1* | 65.94 | 1.47E-10 | 8.09E-08 | *cdca8* | 23.61 | 1.02E-07 | 1.76E-05 |
| *tk1* | 62.11 | 2.48E-08 | 5.76E-06 | *ube2t* | 22.06 | 1.63E-04 | 6.44E-03 |
| *si:dkey-6i22.5* | 59.94 | 2.9E-07 | 4.07E-05 | *ube2c_1* | 21.43 | 9.68E-12 | 7.9E-09 |
| *arhgef39* | 53.80 | 8.26E-10 | 3.19E-07 | *si:ch211-63o20.7* | 21.07 | 2.56E-05 | 1.48E-03 |
| *si:ch211-156j22.4* | 45.29 | 2.39E-06 | 2.2E-04 | *lifrb* | 19.95 | 4.13E-11 | 2.75E-08 |
| *knstrn* | 44.46 | 1.38E-05 | 9.1E-04 | *pane1* | 19.17 | 1.24E-06 | 1.27E-04 |
| *nuf2* | 39.51 | 5.41E-07 | 6.66E-05 | *kpna2* | 19.02 | 5.37E-14 | 9.11E-11 |
| *aurkb* | 38.93 | 2.23E-11 | 1.53E-08 | *rrm2_1* | 18.27 | 2.09E-08 | 5.09E-06 |
| *si:ch211-266i6.3* | 38.08 | 5.4E-06 | 4.22E-04 | *mibp* | 18.10 | 1.29E-04 | 5.33E-03 |
| *top2a* | 36.02 | 5.54E-07 | 6.74E-05 | *nusap1* | 17.95 | 1.14E-06 | 1.19E-04 |
| *prg4a* | 34.49 | 6.61E-12 | 5.6E-09 | *cenpk* | 17.53 | 9.24E-04 | 0.02 |
| *cdk1* | 33.84 | 2.07E-11 | 1.47E-08 | *ccna2* | 17.42 | 1.39E-10 | 7.85E-08 |

Filtering parameters: mean RPKM ≥5, fold change ≥2, false discovery rate (FDR) p-value ≤0.05.

**Table S9.** Full list of Reactome pathways (# of genes ≥5) enriched from 67 upregulated differentially expressed genes at 2dpi compared to 7dpf in macrophage/microglia (mCherry$^+$) cells.

| Reactome pathway | # of genes | Raw p-value |
|---|---|---|
| Cell Cycle, Mitotic | 19 | 5.49E-18 |
| Cell Cycle | 19 | 7E-17 |
| Mitotic Prometaphase | 13 | 3.31E-15 |
| M Phase | 15 | 7.69E-15 |
| Resolution of Sister Chromatid Cohesion | 11 | 2.74E-14 |
| Cell Cycle Checkpoints | 13 | 2.47E-13 |
| Amplification of signal from unattached kinetochores via a MAD2 inhibitory signal | 7 | 6.58E-09 |
| Amplification of signal from the kinetochores | 7 | 6.58E-09 |
| Condensation of Prophase Chromosomes | 5 | 9.96E-09 |
| Cyclin A/B1/B2 associated events during G2/M transition | 5 | 1.2E-08 |
| Mitotic Spindle Checkpoint | 7 | 1.8E-08 |
| RHO GTPase Effectors | 9 | 1.92E-08 |
| RHO GTPases Activate Formins | 7 | 2.9E-08 |
| Separation of Sister Chromatids | 8 | 3.06E-08 |
| Mitotic Anaphase | 8 | 3.76E-08 |
| Mitotic Metaphase and Anaphase | 8 | 3.92E-08 |
| Signaling by Rho GTPases | 10 | 5.77E-08 |
| Mitotic Prophase | 6 | 9.59E-08 |
| Regulation of mitotic cell cycle | 6 | 1.44E-07 |
| APC/C-mediated degradation of cell cycle proteins | 6 | 1.44E-07 |
| G2/M Transition | 7 | 3.56E-07 |
| Mitotic G2-G2/M phases | 7 | 3.85E-07 |
| Activation of APC/C and APC/C:Cdc20 mediated degradation of mitotic proteins | 5 | 2.19E-06 |
| Regulation of APC/C activators between G1/S and early anaphase | 5 | 2.79E-06 |
| Regulation of PLK1 Activity at G2/M Transition | 5 | 4.15E-06 |
| G2/M Checkpoints | 5 | 4.12E-05 |
| Signal Transduction | 15 | 1.9E-04 |
| Innate Immune System | 10 | 4.47E-04 |

**Table S10.** Full list of Reactome pathways (# of genes ≥5) enriched from 208 upregulated differentially expressed genes at 4dpi compared to 9dpf in macrophage/microglia (mCherry$^{+}$) cells.

| Reactome pathway | # of genes | Raw p-value |
|---|---|---|
| Cell Cycle, Mitotic | 55 | 2.63E-43 |
| Cell Cycle | 57 | 1.45E-42 |
| Cell Cycle Checkpoints | 34 | 1.76E-27 |
| S Phase | 26 | 2.53E-24 |
| Activation of ATR in response to replication stress | 18 | 5.12E-24 |
| Synthesis of DNA | 23 | 1.9E-22 |
| DNA Replication | 23 | 7.2E-22 |
| Chromosome Maintenance | 17 | 1.18E-18 |
| Mitotic Prometaphase | 23 | 1.45E-18 |
| DNA strand elongation | 13 | 4.34E-18 |
| Lagging Strand Synthesis | 12 | 3.15E-17 |
| G2/M Checkpoints | 20 | 3.97E-17 |
| PCNA-Dependent Long Patch Base Excision Repair | 12 | 5E-17 |
| Activation of the pre-replicative complex | 13 | 1.27E-16 |
| M Phase | 26 | 1.77E-16 |
| Telomere C-strand (Lagging Strand) Synthesis | 12 | 1.8E-16 |
| Resolution of Abasic Sites (AP sites) | 13 | 2.42E-16 |
| Resolution of AP sites via the multiple-nucleotide patch replacement pathway | 12 | 2.68E-16 |
| Extension of Telomeres | 12 | 3.94E-16 |
| Resolution of Sister Chromatid Cohesion | 17 | 5.25E-15 |
| Base Excision Repair | 13 | 1.97E-14 |
| G1/S Transition | 16 | 2.88E-14 |
| Mitotic G1 phase and G1/S transition | 17 | 3.25E-14 |
| DNA Repair | 22 | 4.43E-14 |
| Telomere Maintenance | 12 | 4.98E-14 |
| Homology Directed Repair | 14 | 5.87E-14 |
| HDR through Homologous Recombination (HRR) or Single Strand Annealing (SSA) | 13 | 4.15E-13 |
| Mitotic Anaphase | 18 | 4.91E-13 |

| Reactome pathway | # of genes | Raw p-value |
|---|---|---|
| Mitotic Metaphase and Anaphase | 18 | 5.35E-13 |
| DNA Double-Strand Break Repair | 15 | 3.73E-12 |
| Gap-filling DNA repair synthesis and ligation in GG-NER | 9 | 4.67E-12 |
| DNA Replication Pre-Initiation | 13 | 5.35E-12 |
| Regulation of TP53 Activity through Phosphorylation | 13 | 6.99E-12 |
| Amplification of signal from unattached kinetochores via a MAD2 inhibitory signal | 13 | 1.17E-11 |
| Amplification of signal from the kinetochores | 13 | 1.17E-11 |
| Translesion synthesis by REV1 | 8 | 1.9E-11 |
| Processive synthesis on the lagging strand | 8 | 1.9E-11 |
| Translesion synthesis by POLI | 8 | 2.83E-11 |
| Translesion synthesis by POLK | 8 | 2.83E-11 |
| Regulation of TP53 Activity | 15 | 3.06E-11 |
| Separation of Sister Chromatids | 16 | 3.5E-11 |
| HDR through Homologous Recombination (HRR) | 11 | 4.01E-11 |
| Recognition of DNA damage by PCNA-containing replication complex | 9 | 4.99E-11 |
| Translesion Synthesis by POLH | 8 | 5.91E-11 |
| Termination of translesion DNA synthesis | 9 | 6.44E-11 |
| Mitotic Spindle Checkpoint | 13 | 6.74E-11 |
| Processive synthesis on the C-strand of the telomere | 7 | 1.43E-10 |
| RHO GTPases Activate Formins | 13 | 1.55E-10 |
| Transcriptional Regulation by TP53 | 18 | 2.01E-10 |
| Processing of DNA double-strand break ends | 9 | 2.57E-10 |
| Presynaptic phase of homologous DNA pairing and strand exchange | 9 | 3.17E-10 |
| Translesion synthesis by Y family DNA polymerases bypasses lesions on DNA template | 9 | 3.17E-10 |
| Polymerase switching | 7 | 3.44E-10 |
| Polymerase switching on the C-strand of the telomere | 7 | 3.44E-10 |
| Leading Strand Synthesis | 7 | 3.44E-10 |
| Removal of the Flap Intermediate | 7 | 5.12E-10 |
| Homologous DNA Pairing and Strand Exchange | 9 | 5.76E-10 |

| Reactome pathway | # of genes | Raw p-value |
| --- | --- | --- |
| Dual Incision in GG-NER | 9 | 5.76E-10 |
| Global Genome Nucleotide Excision Repair (GG-NER) | 11 | 6.57E-10 |
| Nucleotide Excision Repair | 12 | 1.04E-09 |
| Gap-filling DNA repair synthesis and ligation in TC-NER | 10 | 1.28E-09 |
| DNA Damage Bypass | 9 | 1.7E-09 |
| G2/M DNA damage checkpoint | 10 | 3.2E-09 |
| HDR through Single Strand Annealing (SSA) | 8 | 3.21E-09 |
| Orc1 removal from chromatin | 10 | 3.62E-09 |
| Removal of the Flap Intermediate from the C-strand | 6 | 4.08E-09 |
| Transcription-Coupled Nucleotide Excision Repair (TC-NER) | 10 | 5.19E-09 |
| RHO GTPase Effectors | 15 | 1.7E-08 |
| Mismatch repair (MMR) directed by MSH2:MSH6 (MutSalpha) | 6 | 1.92E-08 |
| Dual incision in TC-NER | 9 | 2.25E-08 |
| Switching of origins to a post-replicative state | 10 | 2.6E-08 |
| Mismatch Repair | 6 | 2.66E-08 |
| Condensation of Prometaphase Chromosomes | 5 | 7.45E-08 |
| Mismatch repair (MMR) directed by MSH2:MSH3 (MutSbeta) | 5 | 1.71E-07 |
| TP53 Regulates Transcription of Cell Cycle Genes | 7 | 2.26E-07 |
| Generic Transcription Pathway | 21 | 2.63E-07 |
| Assembly of the pre-replicative complex | 8 | 4.81E-07 |
| Signaling by Rho GTPases | 16 | 6.6E-07 |
| Regulation of mitotic cell cycle | 8 | 2.27E-06 |
| APC/C-mediated degradation of cell cycle proteins | 8 | 2.27E-06 |
| RNA Polymerase II Transcription | 21 | 3.33E-06 |
| SUMOylation of DNA replication proteins | 6 | 4.9E-06 |
| G2/M Transition | 10 | 5.47E-06 |
| Mitotic G2-G2/M phases | 10 | 6.05E-06 |
| Cyclin A/B1/B2 associated events during G2/M transition | 5 | 6.1E-06 |
| Gene expression (Transcription) | 22 | 8.48E-06 |

| | | |
|---|---|---|
| Nucleosome assembly | 5 | 2.74E-05 |
| Deposition of new CENPA-containing nucleosomes at the centromere | 5 | 2.74E-05 |
| Fanconi Anemia Pathway | 5 | 3.96E-05 |
| Metabolism of nucleotides | 7 | 4.33E-05 |
| AURKA Activation by TPX2 | 6 | 8.16E-05 |
| SUMO E3 ligases SUMOylate target proteins | 8 | 1.08E-04 |
| Regulation of APC/C activators between G1/S and early anaphase | 6 | 1.32E-04 |
| SUMOylation | 8 | 1.4E-04 |
| Cellular responses to stress | 12 | 2.73E-04 |
| Cellular responses to external stimuli | 12 | 2.79E-04 |
| Signal Transduction | 35 | 3.6E-04 |
| Activation of APC/C and APC/C:Cdc20 mediated degradation of mitotic proteins | 5 | 8.45E-04 |
| Mitotic Prophase | 5 | 1.05E-03 |
| Innate Immune System | 21 | 1.45E-03 |
| Regulation of PLK1 Activity at G2/M Transition | 5 | 1.51E-03 |
| Regulation of Complement cascade | 5 | 2.6E-03 |
| Immune System | 29 | 3.08E-03 |
